# *In situ* imaging of bacterial membrane projections and associated protein complexes using electron cryo-tomography

**DOI:** 10.1101/2021.07.13.452161

**Authors:** Mohammed Kaplan, Georges Chreifi, Lauren Ann Metskas, Janine Liedtke, Cecily R Wood, Catherine M Oikonomou, William J Nicolas, Poorna Subramanian, Lori A Zacharoff, Yuhang Wang, Yi-Wei Chang, Morgan Beeby, Megan Dobro, Mark J McBride, Ariane Briegel, Carrie Shaffer, Grant J Jensen

**Affiliations:** Division of Biology and Biological Engineering, California Institute of Technology, Pasadena, CA 91125, USA; Leiden University, Sylvius Laboratories, Leiden, The Netherlands; University of Kentucky Department of Veterinary Science; University of Southern California, Department of Physics and Astronomy, Los Angeles, CA90089; Department of Biochemistry and Biophysics, Perelman School of Medicine, University of Pennsylvania, Philadelphia, PA 19104, USA; Department of Life Sciences, Imperial College London, South Kensington Campus, London SW7 2AZ, UK; Hampshire College, Amherst, Massachusetts, USA; Department of Biological Sciences, University of Wisconsin-Milwaukee, Milwaukee, Wisconsin, USA; University of Kentucky Department of Microbiology, Immunology, and Molecular Genetics; University of Kentucky Department of Pharmaceutical Sciences; Department of Chemistry and Biochemistry, Brigham Young University, Provo, UT 84604, USA

## Abstract

The ability to produce membrane projections in the form of tubular membrane extensions (MEs) and membrane vesicles (MVs) is a widespread phenomenon among bacteria. Despite this, our knowledge of the ultrastructure of these extensions and their associated protein complexes remains limited. Here, we surveyed the ultrastructure and formation of MEs and MVs, and their associated protein complexes, in tens of thousands of electron cryo-tomograms of ∼ 90 bacterial species that we have collected for various projects over the past 15 years (Jensen lab database), in addition to data generated in the Briegel lab. We identified MEs and MVs in 13 species and classified several major ultrastructures: 1) tubes with a uniform diameter (with or without an internal scaffold), 2) tubes with irregular diameter, 3) tubes with a vesicular dilation at their tip, 4) pearling tubes, 5) connected chains of vesicles (with or without neck-like connectors), 6) budding vesicles and nanopods. We also identified several protein complexes associated with these MEs and MVs which were distributed either randomly or exclusively at the tip. These complexes include a secretin-like structure and a novel crown-shaped structure observed primarily in vesicles from lysed cells. In total, this work helps to characterize the diversity of bacterial membrane projections and lays the groundwork for future research in this field.

## Introduction

Membrane extensions and vesicles (henceforth referred to as MEs and MVs) have been described in many types of bacteria. They are best characterized in diderms, where they stem mainly from the outer membrane (OM; we thus refer to OMEs and OMVs) and perform a variety of functions [1–4]. For example, the OMEs of *Shewanella oneidensis* (aka nanowires) are involved in extracellular electron transfer [5, 6]. The OM tubes of *Myxococcus xanthus* are involved in the intra-species transfer of periplasmic and OM-associated material between different cells that is essential for the complex social behavior of this species [7–9]. The OMVs of *Vibrio cholerae* act as a defense mechanism, helping the bacterium circumvent phage infection [10]. A marine Flavobacterium affiliated with the genus *Formosa* (strain Hel3_A1_48) extrudes membrane tubes and vesicles that contain the type IX secretion system and digestive enzymes [11]. OMVs often function in pathogenesis. The OM blebs and vesicles of *Flavobacterium psychrophilum* have proteolytic activities that help release nutrients from the environment and impede the host immune system [12]. The OMVs of *Francisella novicida* contain virulence factors, suggesting they are involved in pathogenesis [13]. Similarly, the virulence of *Flavobacterium columnare* is associated with the secretion of OMVs [14], and membrane tubes and secreted vesicles have been observed in other, human pathogens like *Helicobacter pylori* and *Vibrio vulnificus* [15, 16].

MEs and MVs are also produced by monoderm bacteria and archaea. MVs stemming from the cytoplasmic membrane of Gram-positive bacteria have been reported to encapsulate DNA (see Ref. [17] and references therein.) Membrane nanotubes were recently discovered in the Gram-positive *Bacillus subtilis*, as well as the Gram-negative *Escherichia coli.* These nanotubes were found to connect two different bacterial cells and are involved in the transfer of cytoplasmic material between bacterial cells of the same and different species, and even to eukaryotic cells [18–24].

The structures of MEs and MVs are as varied as their functions. While *S. oneidensis* nanowires are chains of interconnected outer membrane vesicles with variable diameter and decorated with cytochromes [6], OM tubes of *H. pylori* have a fixed diameter of ∼40 nm and are characterized by an inner scaffold and lateral ports [15]. *V. vulnificus* produces tubes from which vesicles ultimately pinch off by biopearling, forming a regular concentric pattern surrounding the cell [16]. Cells with an external surface layer (S-layer) can produce structures known as “nanopods,” which consist of membrane vesicles inside a sheath of S-layer. These have been reported in the soil-residing bacterium *Delftia* sp. Cs1-4 [25] and archaea of the order *Thermococcales* [26]. Finally, some diderms produce DNA-containing MVs consisting of both inner and outer membranes (see Ref. [4] and references therein).

Different models have been proposed for how MEs and MVs form. In diderms, membrane blebbing may occur due to changes in the periplasmic turgor pressure, lipopolysaccharide repulsion or alterations in the contacts between the OM and the peptidoglycan cell wall [4]. Chains of interconnected vesicles are often observed, either as a result of direct vesicular budding from the OM or due to biopearling of membrane tubes [6, 11]. Formation of tubes is thought to be a stabilizing factor as it results in smaller vesicles, with tubes pearling into distal chains of vesicles that eventually disconnect [27]. Other extensions may be formed by dedicated machinery. Interestingly, nanotubes involved in cytoplasmic exchange have been reported to be dependent on a conserved set of proteins involved in assembly of the flagellar motor known as the type III secretion system core complex (CORE): FliP/O/Q/R and FlhA/B [18, 24]. Recently, it was also shown that the formation of bacterial nanotubes significantly increases under stress conditions or in dying cells, caused by biophysical forces resulting from the action of the cell wall hydrolases LytE and LytF [28].

Structural studies of MEs and MVs have relied mainly on scanning electron microscopy (SEM), conventional transmission electron microscopy (TEM), and light (fluorescence) microscopy. While these methods have significantly advanced our understanding, they are limited in terms of the information they can provide. For instance, in SEM and conventional TEM, sample preparation such as fixation, dehydration, and staining disrupt membrane ultrastructure. While light microscopy can reveal important information about the dynamics and timescales on which MEs and MVs form (e.g. [29]), no ultrastructural details can be resolved; MEs and MVs of different morphology appear identical. Currently, only electron cryo-tomography (cryo-ET) allows visualization of structures in a near-native state inside intact (frozen-hydrated) cells with macromolecular (∼5 nm) resolution. This method has already been invaluable in revealing the structures of several membrane extensions, including *S. oneidensis* nanowires [6], *H. pylori* nanotubes [15], *D. acidovorans* nanopods [25], *V. vulnificus* OMV chains [16], and more recently cell-cell bridges in the archaeon *Haloferax volcanii* [30].

To understand what membrane extensions exist in bacterial cells and how they might form, we undertook a survey of ∼90 bacterial species, drawing on a database of tens of thousands of electron cryo-tomograms of intact cells collected by our group for various projects over the past 15 years [31, 32], in addition to data generated in the Briegel lab. Our survey revealed membrane projections in 13 bacterial species. These projections took various forms: 1) tubes with a uniform diameter and with an internal scaffold, 2) tubes with a uniform diameter and without a clear internal scaffold, 3) tubes with a vesicular dilation at their tip (teardrop-like extensions), 4) tubes with irregular diameter or pearling tubes, 5) interconnected chains of vesicles with uniform neck-like connectors, budding or detached OMVs, and 7) nanopods. We also identified protein complexes associated with MEs and MVs in these species. These complexes were either randomly distributed on the MEs and MVs or exhibited a preferred localization at their tip.

## Results

We examined tens of thousands of electron cryo-tomograms of ∼ 90 bacterial species collected in the Jensen lab for various projects over the past 15 years together with tomograms collected in the Briegel lab. Most cells were intact, but some had naturally lysed. Note that we make this classification based on the cells’ appearance in tomograms; intact cells have an unbroken cell envelope, uniform periplasmic width, and consistently dense cytoplasm. In addition to cryo-tomograms of cells, this dataset also included naturally-shed vesicles purified from *S oneidensis*. In all, we identified OMEs and OMVs in 13 bacterial species (summarized in Table S1).

### I- The diverse forms of bacterial membrane structures

Based on their features, we classified membrane projections into the following categories: **1)** tubular extensions with a uniform diameter and with an internal scaffold (Fig. 1 a & b); **2)** tubular extensions with a uniform diameter and without a clear internal scaffold (Fig. 1 c-g ); **3)** tubular extensions with a vesicular dilation at the tip (a teardrop-like structure) and irregular dark densities inside (Fig. 1h); **4)** tubular extensions with irregular diameter or pearling tubes (Fig. 2 a-g); **5)** interconnected chains of vesicles with uniform neck-like connectors (Fig. 2 h & i); **6)** budding or detached vesicles: budding vesicles were still attached to the membrane, while detached vesicles were observed near a cell and could have budded directly or from a tube that pearled (Fig. 3 a-d); nanopods: tubes of S-layer containing OMVs (Fig. 3 e-i). See Table S1 for a summary of these observations.

**Figure 1:**
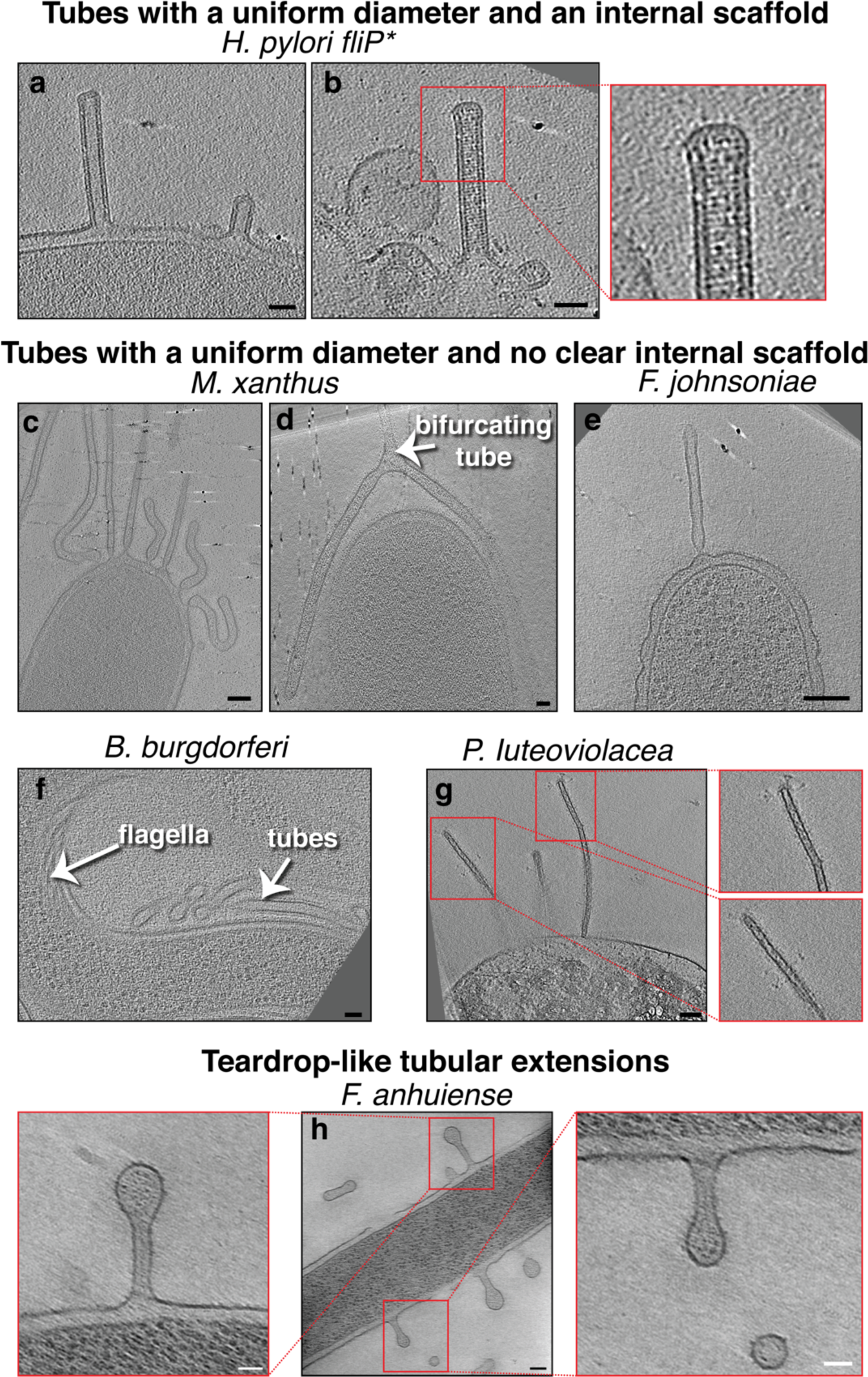
Membrane tubes with a uniform diameter, either with or without an internal scaffold. Slices through electron cryo-tomograms of the indicated bacterial species highlighting the presence of OMEs with uniform diameters and either with **(a-b)** or without **(c-g)** an internal scaffold, and teardrop-like extensions (**h**). In this and all subsequent figures, red boxes indicate enlarged views of the same slice. Scale bars are 50 nm, except in main panel (h) 100 nm.

**Figure 2:**
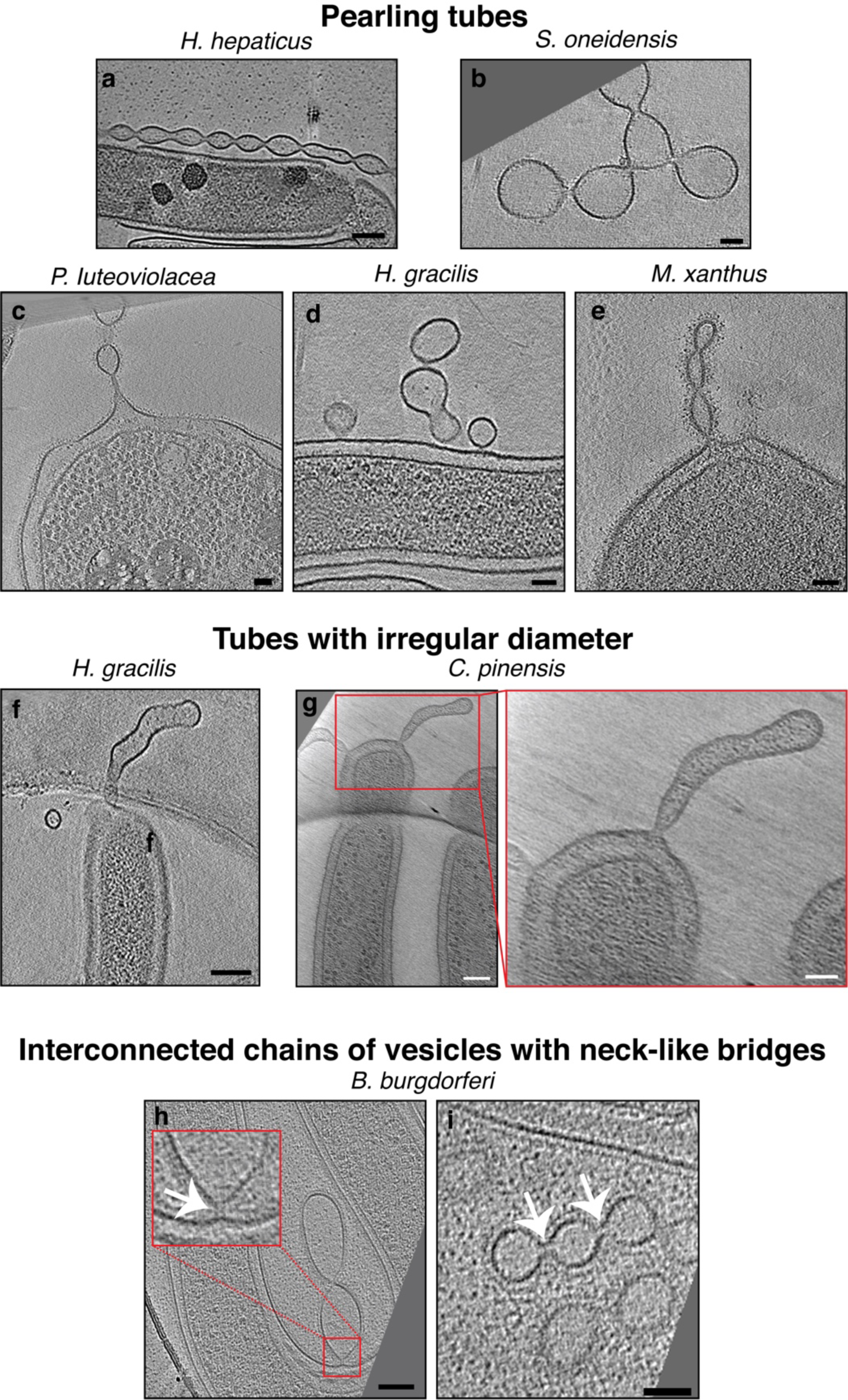
Pearling tubes, tubes with irregular diameter and vesicle chains with neck-like connections. Slices through electron cryo-tomograms of the indicated bacterial species highlighting the presence of pearling tubes **(a-e)**, tubes with irregular diameter **(f-g)**, or OMV chains connected by neck-like bridges **(h-i)**. White arrows in the enlargement in **(h)** and in panel **(i)** point to the 14-nm connectors in *B. burgdorferi*. Scale bars are 50 nm, except in main panel (g) 100 nm.

**Figure 3:**
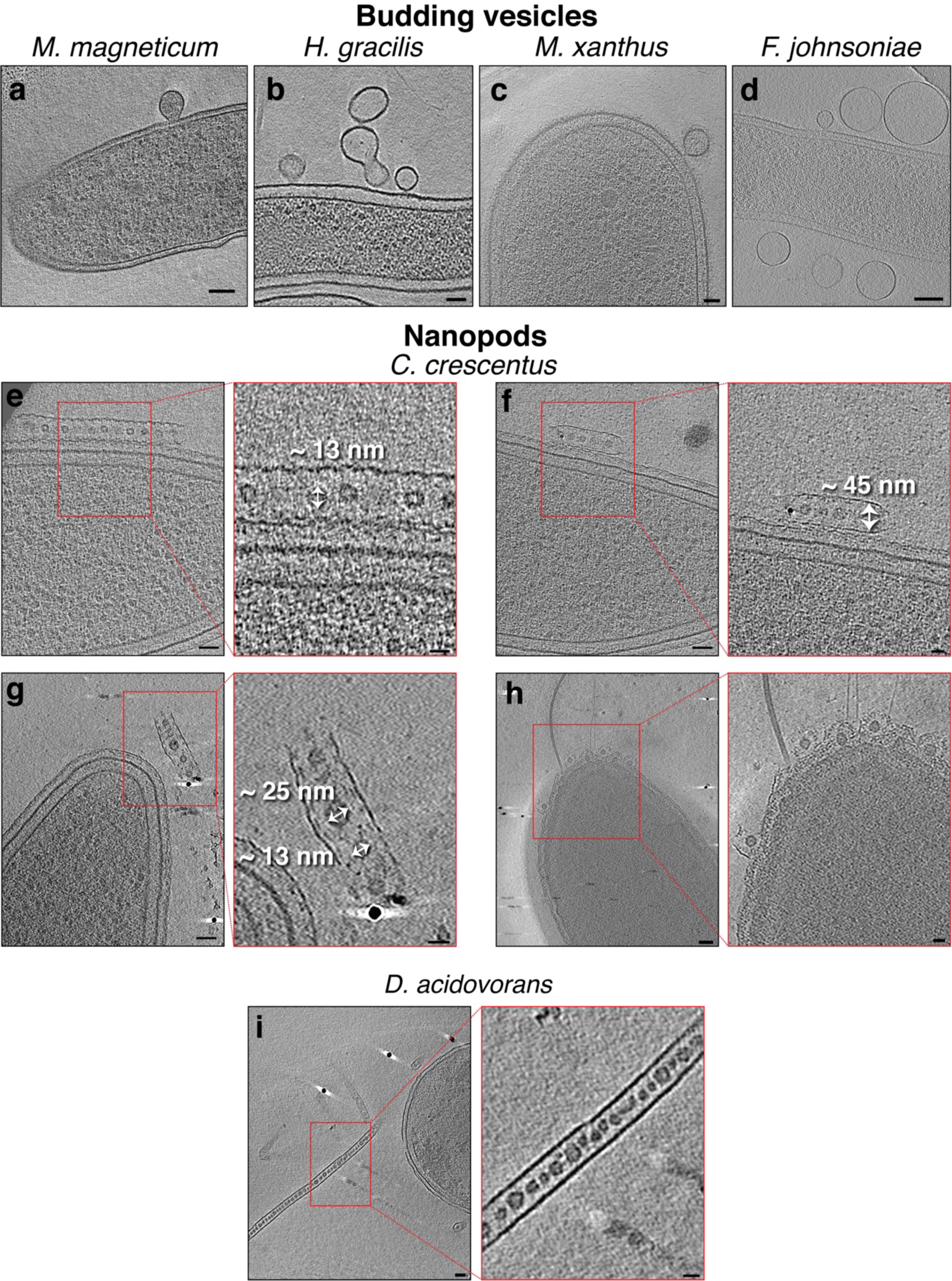
Budding OMVs and nanopods. Slices through electron cryo-tomograms of the indicated bacterial species highlighting the presence of budding vesicles **(a-d)** or nanopods **(e- i)**. Scale bars are 50 nm in main panels and 20 nm in enlargements.

Scaffolded membrane tubes were observed only in *H. pylori* and had a uniform diameter of 40 nm. The *H. pylori* strain imaged (*fliP\**) contains a naturally-occurring point mutation that disrupts the function of FliP, the platform upon which other CORE proteins assemble [33–35]. In addition, the dataset contained other mutants in this *fliP\** background including additional CORE proteins (Δ*fliO* and Δ*fliQ*), flagellar basal body proteins (Δ*fliM* and Δ*fliG*), and the tyrosine kinase required for expression of the class II flagellar genes (Δ*flgS*) [36] (Figs. 1 a-b and 4). This suggests that the *H. pylori* membrane tubes are unrelated to the CORE-dependent nanotubes that mediate cytoplasmic exchange in *B. subtilis* and other species [18, 24].

Previously, *H. pylori* tubes were described as forming in the presence of eukaryotic host cells [15]. Here, however, we observed tubes on *H. pylori* grown on agar plates in the absence of eukaryotic cells, suggesting that they also form in the absence of host cells. We observed some differences, though, from the tubes formed in the presence of host cells: the tube ends were closed, no clear lateral ports were seen, and the tubes were usually straight. While some of these tubes extended more than 0.5 μm, we never observed pearling. However, in some tubes, the internal scaffold did not extend all the way to the tip, and its absence caused the tube to dilate (from 40 nm in the presence of the scaffold to 66 nm in its absence, see Fig. 4f). In some cases we also observed tubes stemming from vesicles resulting from cell lysis (Figs. 4f and S1).

**Figure 4:**
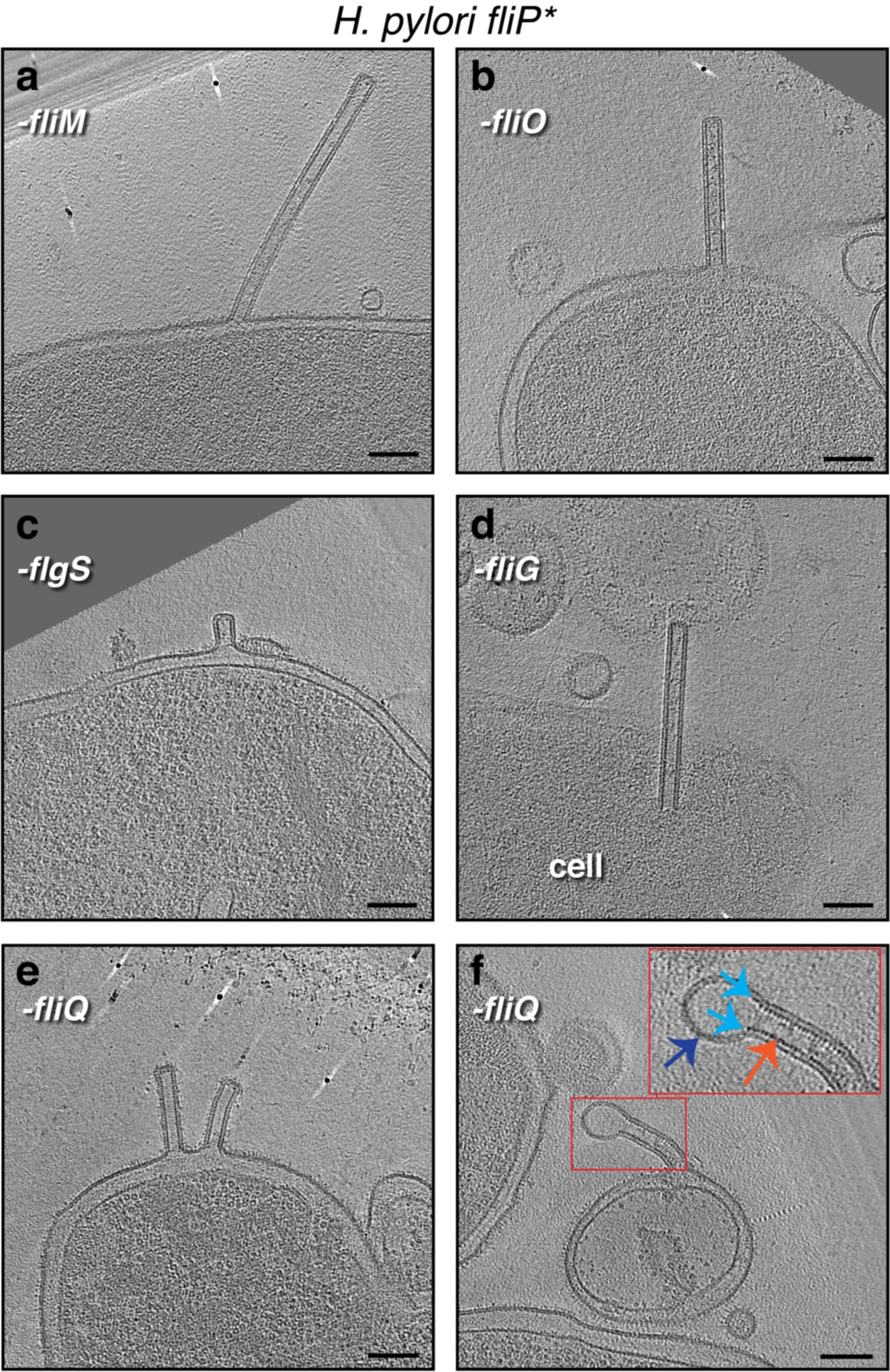
The formation of OM tubes persists in various *H. pylori* mutants, including CORE mutants. Slices through electron cryo-tomograms of the indicated *H. pylori* mutants (all in the *fliP\** background) showing the presence of membrane tubes. The enlargement in **(f)** highlights a dilation at the end of the tube (dark blue arrow) due to the absence of the scaffold (orange arrow). Light blue arrows indicate the end points of the scaffold. Scale bars are 100 nm.

In *Flavobacterium anhuiense* and *Chitinophaga pinensis*, which are both endophytic species extracted from sugar beet roots, in addition to tubes with irregular diameter and OMVs, tubular extensions with a uniform diameter and a vesicular dilation (teardrop-like structure) were observed stemming from the sides of the cell (Fig. 1h). Interestingly, irregular dark densities were observed inside these teardrop-like extensions (Fig. 1h). Chains of vesicles connected by neck-like bridges were similarly observed in a single species: *Borrelia burgdorferi*. The bridges were consistently ∼14 nm in length and ∼8 nm in width. Where chains were seen attached to the outer membrane, a neck-like connection was present at the budding site (Fig. 2h). Vesicles in each chain were of a uniform size, usually 35-40 nm wide (e.g. Fig. 2i), but occasionally larger (e.g. Fig. 2h).

When both tubes and vesicles were observed in the same species, the tubes generally had a more uniform diameter than the vesicles, which were of variable sizes and often had larger diameters than the tubes (Figs. S2 & S3). In addition, when a tube pearled into vesicles, there was no clear correlation between the length of the tube and the initiation point of pearling, with some tubes extending for many micrometers without pearling while other, shorter tubes were in the process of forming vesicles (Movies S1, S2, S3 and Fig. 2e). While most pearling was seen at the tips of tubes, pearling occasionally occurred simultaneously at both proximal and distal ends of the same tube (Movie S3). With one exception, pearling was seen in all species with tubes of uniform diameter and no internal scaffold. The exception was lysed *Pseudoalteromonas luteoviolacea*, which had narrow tubes only 20 nm in diameter (Fig. 1g). Some lysed *P. luteoviolacea* contained wider, pearling tubes (Fig. 2c). Interestingly, the tubes of various *M. xanthus* strains (see Materials and Methods) and *P. luteoviolacea* could bifurcate into branches, each of which had a uniform diameter similar to that of the main branch (Movie S4 and Fig. 1d and Fig. S4).

In *C. crescentus* tomograms, we identified structures very similar to the “nanopod” extensions previously reported in *D. acidovorans* [25]. These structures consist of a tube made of the S-layer encasing equally-spaced OMVs (Fig. 3 e-h and Movie S5). The diameter of the S-layer tubes was ∼45 nm and vesicles exhibited diameters ranging from ∼13-25 nm. The nanopods were seen either detached from the cell (Fig. 3 e-g), or budding from the pole of *C. crescentus* (Fig. 3h).

### II- Protein complexes associated with membrane structures

Next, we examined protein complexes associated with OMEs and OMVs that we could identify in our cryo-tomograms. These complexes fell into three categories: **1)** randomly-located complexes found on OMEs, OMVs and cells; **2)** randomly-located complexes observed only on OMEs and OMVs, and **3)** complexes exclusively located at the tip of OMEs/OMVs.

In the first category, we observed what appeared to be the OM-associated portion of the empty basal body of the type IVa pilus (T4aP) machinery in OMEs of *M. xanthus.* These complexes, which were also found in the OM of intact cells, did not exhibit a preferred localization within the tube (Fig. 5a & b).

**Figure 5:**
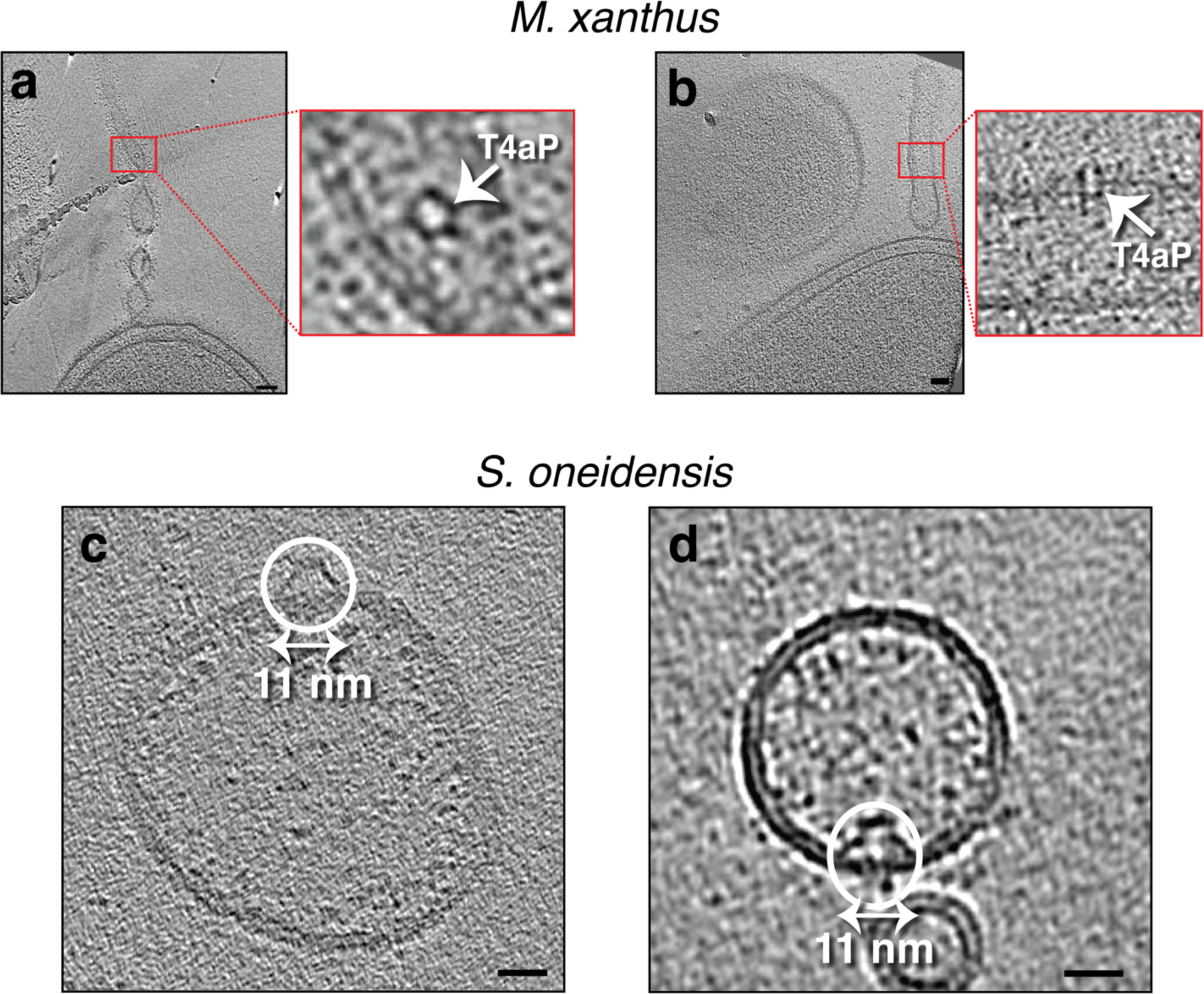
Randomly-located protein complexes on OMEs of *M. xanthus* and purified MVs of *S. oneidensis*. a & b) Slices through electron cryo-tomograms of *M. xanthus* indicating the presence of pearling tubes with top (a) and side (b) views of type IVa pilus basal bodies (T4aP). Scale bars are 50 nm. c & d) Slices through electron cryo-tomograms of purified *S. oneidensis* naturally-shed MEs and MVs highlighting the presence of trapezoidal structures on the outside (c) and inside (d) of vesicles. Scale bars are 10 nm.

The second category of protein complexes, observed only on MEs and not on cells, contained two structures. The first was a trapezoidal structure observed on purified OMVs of *S. oneidensis*. The structure was ∼11 nm wide at its base at the membrane and was seen sometimes on the outside (Fig. 5c) and sometimes the inside of vesicles (Fig. 5 d). The second structure was a large crown-like complex. We first observed these complexes on the outer surface of membrane vesicles associated with lysed *M. xanthus* cells (Fig. 6a). Occasionally, they were also present on what appeared to be the inner leaflet of the inner membrane of lysed cells (Fig. 6b). The exact topology is difficult to determine, however, since the arrangement of inner and outer membranes can be confounded by cell lysis. The structure of this complex was consistent enough to produce a subtomogram average from nine examples, improving the signal-to-noise ratio and revealing greater detail (Fig. 6c). These crown-like complexes are ∼40 nm tall with a concave top and a base ∼35 nm wide at the membrane (Fig. 6c). No such complexes were seen on OMEs and OMVs associated with intact *M. xanthus* cells. We identified a morphologically similar crown-like complex on the outside of some tubes and vesicles purified from *S. oneidensis* (Fig. 6d-f). However, this complex was smaller, ∼15 nm tall and ∼20 nm wide at its base. As these were purified OMEs/OMVs, we cannot know whether they stemmed from lysed or intact cells. Interestingly, we found a similar large crown-like structure associated with lysed cells of two other species in which we did not identify MEs, namely *Pseudomonas flexibilis* and *Pseudomonas aeruginosa* (Fig. 6g-j & S5).

**Figure 6:**
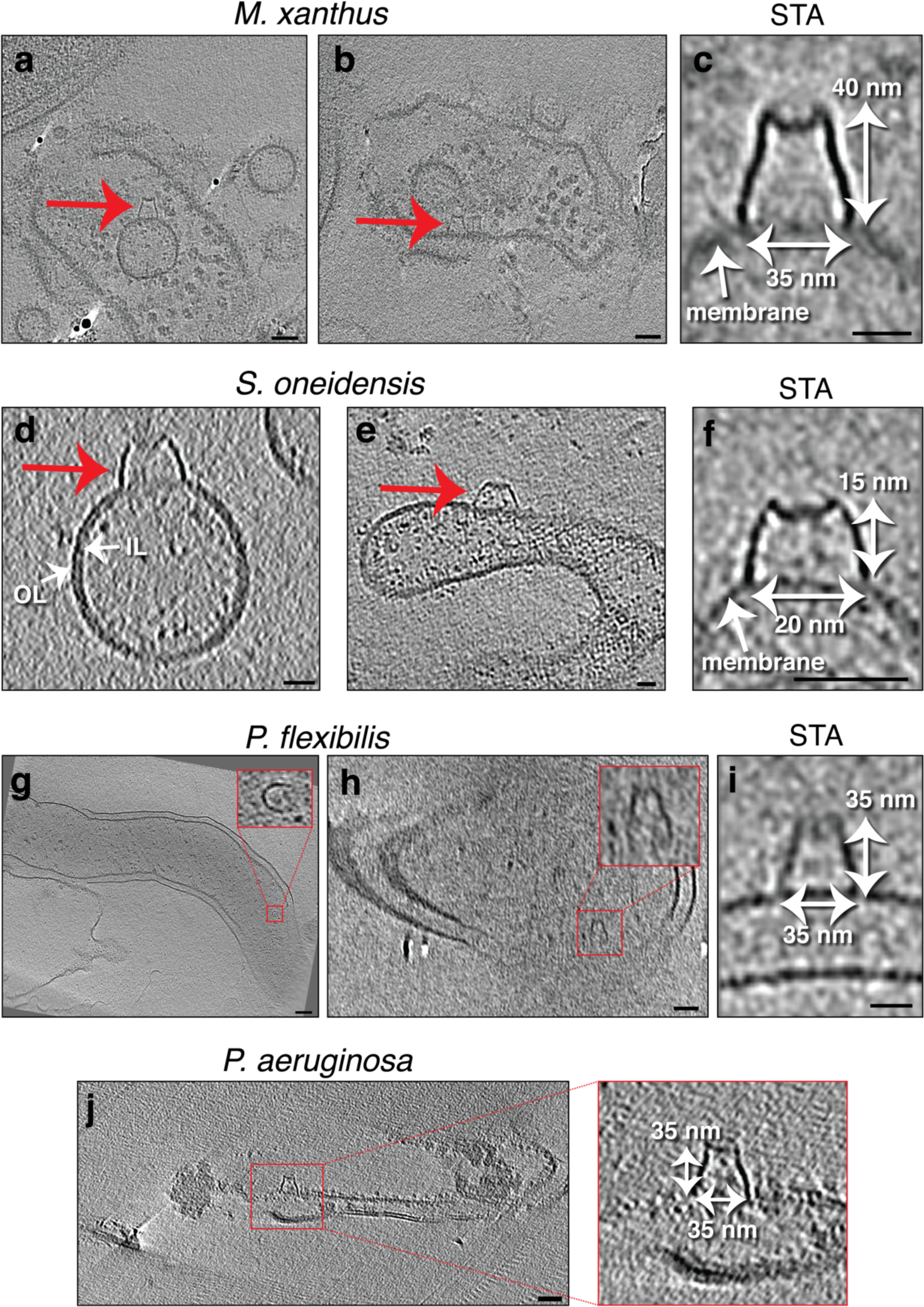
Randomly-located protein complexes associated with lysed cells. Slices through electron cryo-tomograms of lysed cells **(a, b, g, h &j)** or purified MEs and MVs **(d &e)** showing the presence of membrane vesicles and lysed membranes with a crown-like complex (red arrows and red-boxed enlargements). Scale bars: 50 nm (a, b, h and j), 100 nm (g), 10 nm (d and e). **c, f & i)** Central slices through subtomogram averages (with two-fold symmetry along the Y-axis applied) of 9 particles (c), 4 particles (f) or 3 particles (i) of the crown-like complex in the indicated species. Scale bars are 20 nm. OL=outer leaflet, IL=inner leaflet.

In the third category, we observed a secretin-like complex in many tubes and vesicles of *F. johnsoniae*. In tubes attached to the cell, the complex was always located at the distal tip (Figs. 7 & S6 and Movie S6). From 35 membrane tubes seen attached to cells, we identified a secretin-like complex at the tip of 25 of them (∼70%). In OMEs disconnected from the cell, the secretin-like complex was always located at one end (Fig. 7b & e). In total, in 198 tomograms we identified 88 secretin-like particles, none of which were located in the middle of a tube. As the MEs are less crowded than cellular periplasm and usually thinner than intact cells, we could clearly distinguish an extracellular density and three periplasmic densities in side views (red and purple arrows, respectively, in Fig. 7a). Top views showed a plug in the center of the upper part of the complex (yellow arrows in Fig. 7g & h). Subtomogram averaging revealed details of the complex, including the plug and a distinct lower periplasmic ring (Fig. 7i & j & S7). While the upper two periplasmic rings were clearly distinguishable in many of the individual particles (e.g. Fig. 7a), they did not resolve as individual densities in the subtomogram average (Fig. 7i). The extracellular density was not resolved at all in the average, suggesting flexibility in this part.

**Figure 7:**
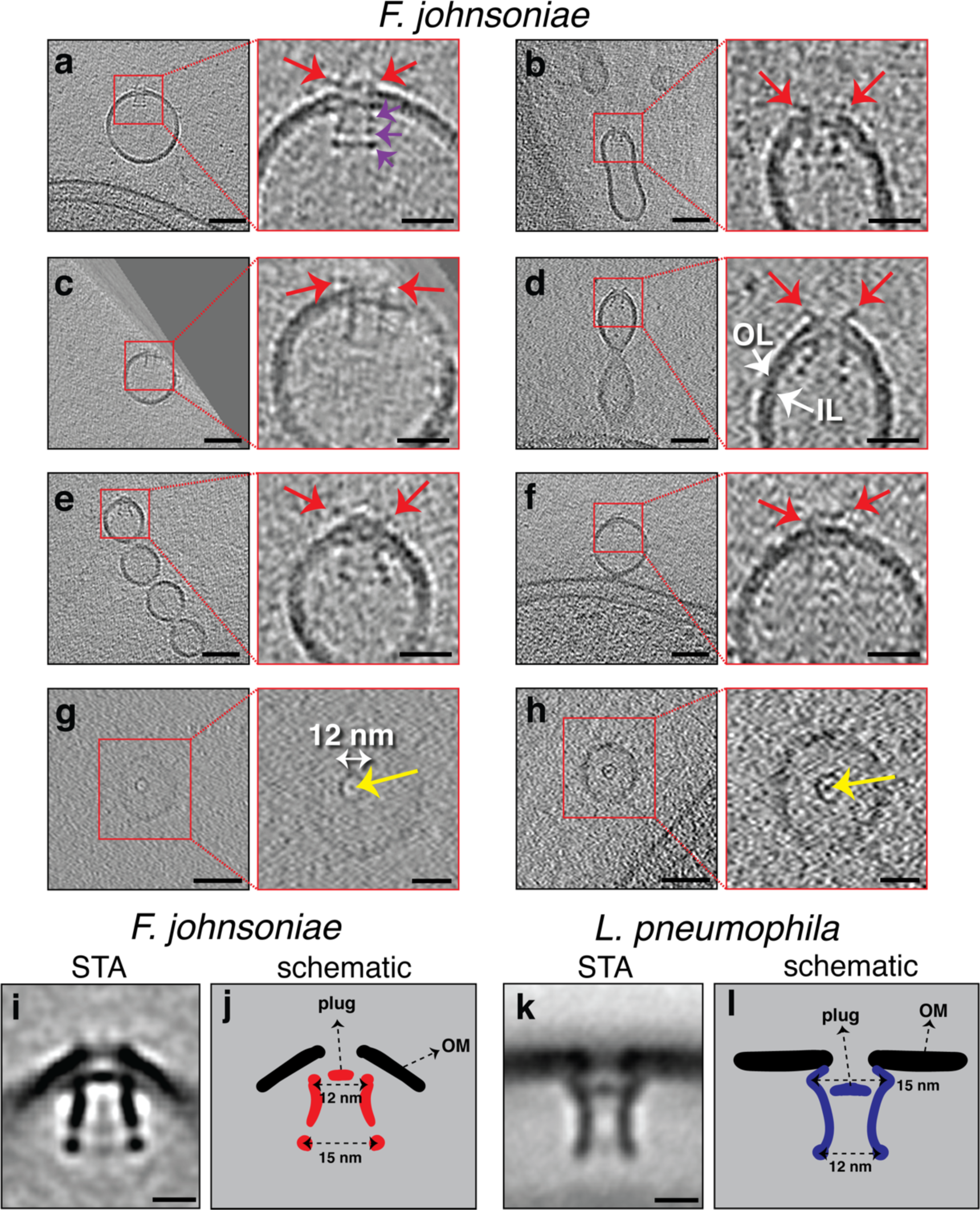
Secretin-like complexes located at the tip of OMEs and OMVs in *F. johnsoniae*. Slices through electron cryo-tomograms of *F. johnsoniae* illustrating the presence of secretin-like complexes (side views in **a-f**, top views in **g & h** with yellows arrows pointing to the plug) in OMEs and OMVs of *F. johnsoniae*. Red arrows point to the extracellular part of the complex. Purple arrows in the enlargement in (a) point to the three periplasmic densities. Scale bars are 50 nm in main panels and 20 nm in enlargements. **i)** A central slice through the subtomogram average of 88 particles of the secretin-like complex (with two-fold symmetry along the Y-axis applied). Scale bar is 10 nm. **j)** A schematic representation of the STA shown in (i). **k)** A central slice through the subtomogram average of the secretin of the T2SS of *L. pneumophila* (EMD 20713, see Ref. [38]). Scale bar is 10 nm. **l)** A schematic representation of the STA shown in (k).

Previous studies showed that a species which belongs to the same phylum as *F. johnsoniae*, namely *Cytophaga hutchinsonii*, uses a putative T2SS to degrade cellulose [37]. Since *F. johnsoniae* also degrades polysaccharides and other polymers, we BLASTed the sequence of the well-characterized *V. cholerae* T2SS secretin protein, GspD (UniProt ID P45779), against the genome of *F. johnsoniae* and found a hit, GspD-like T2SS secretin protein (A5FMB4), with an e-value of 1e^-9^. This result and the general morphological similarity of this secretin to the published structure of the T2SS [38] suggested that the complex we observed might be the secretin of a T2SS. We therefore compared our subtomogram average with the only available *in situ* structure of a T2SS, a recent subtomogram average of the *Legionella pneumophila* T2SS [38] (Fig. 7i-l). The two structures were generally similar in length and both had a plug in the upper part of the complex. However, we also observed differences between the two structures. In *L. pneumophila*, the widest part of the secretin (15 nm) is located near the plug close to the OM, and the lower end of the complex is narrower (12 nm). In *F. johnsoniae*, this topology is reversed, with the narrowest part near the plug and OM (Fig. 7i-l). Additionally, the lowest domain of the *L. pneumophila* secretin did not resolve into a distinct ring as we saw in *F. johnsoniae* and no extracellular density was observed in *L. pneumophila*, either in the subtomogram average or single particles [38].

## Discussion

Our results highlight the diversity of MEs and MVs structures that bacteria can form even within a single species (Fig. 8). For example, we saw two types of membrane tubes in lysed *P. luteoviolacea* cells: one narrower with a uniform diameter of 20 nm which did not pearl into vesicles, and one wider with a variable diameter that did pearl into vesicles (Fig. 1g and Fig. 2c), a distinction which suggests that these extensions play different roles. Similarly, interspecies differences likely reflect different functions. For instance, the tubes of *M. xanthus* were on average longer, more abundant and more branched than the MEs of other species (Movies S1 & S4), which is likely related to their role in communication between cells of this highly social species. However, one interesting observation in all the species we investigated here is that there was no clear distinctive molecular machine at the base of the membrane projections, raising the question of what drives their formation. This observation is consistent with a recent study which showed that liquid-like assemblies of proteins in membranes can lead to the formation of tubular extensions without the need for solid scaffolds [39].

**Figure 8:**
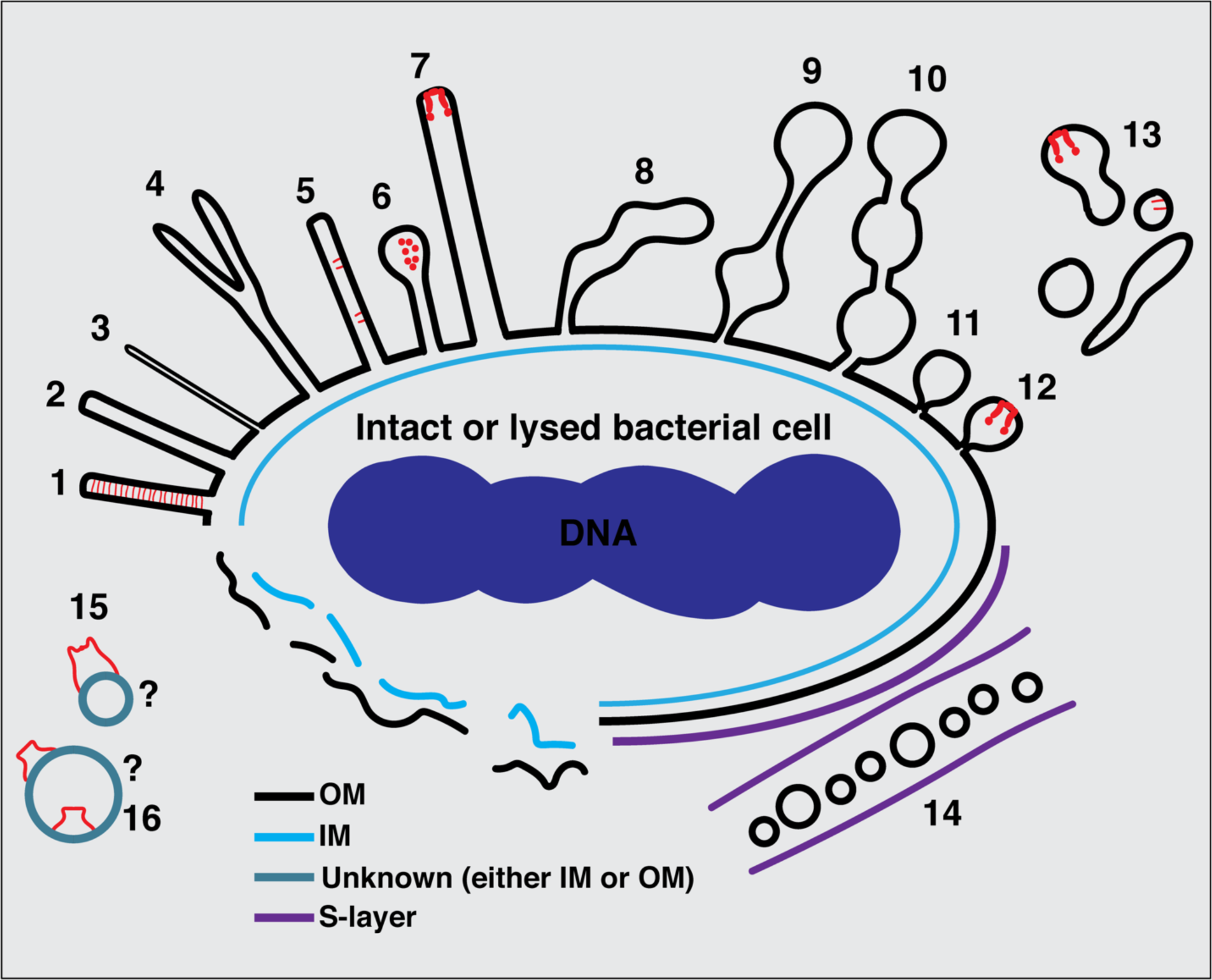
Summary of types of MEs and MVs identified in this study. **1**) tubes with a uniform diameter and with an internal scaffold; **2 & 3**) tubes with a uniform diameter but without an internal scaffold; **4**) bifurcating tubes; **5**) tubes with randomly-located protein complexes (T4aP); **6**) teardrop-like extensions; **7**) tubes with a secretin-like complex at their tip; **8**) tubes with irregular diameter; **9**) pearling tubes; **10**) interconnected chains of vesicles with 14-nm connectors; **11**) budding vesicles; **12**) budding vesicles with a secretin-like complex at their tip; **13**) various disconnected membrane structures in the vicinity of bacterial cells; **14**) nanopods in species with an inner membrane (IM), outer membrane (OM), and S-layer; **15**) membrane structures with a crown-like complex from lysed cells; **16**) purified OMVs with trapezoidal complexes. The question marks in (15) and (16) indicate the difficulty of determining whether a membrane structure from lysed cells or purified vesicles originated from the IM or the OM or is in its original topology.

The scaffolded uniform tubes of *H. pylori* that we observed were formed in samples not incubated with eukaryotic cells, indicating that they can also form in their absence. However, the tubes we found had closed ends and no clear lateral ports, while some of the previously-reported tubes (formed in the presence of eukaryotic host cells) had open ends and prominent ports [15]. It is possible that such features are formed only when *H. pylori* are in the vicinity of host cells. We also show that the tubes of *H. pylori* are CORE-independent, indicating that they are different from the CORE-dependent nanotubes described in other species.

A recent study showed that the formation of bacterial tubes significantly increases when cells are stressed or dying [28]. Consistent with this, in our cryo-tomograms we saw many MEs and MVs associated with lysed cells (such as in *H. pylori*, *H. hepaticus*, and *P. luteoviolacea*). We also saw tubes and vesicles stemming from intact cells. Given the nature of cryo-ET snapshots, we cannot tell whether a cell that appears intact is stressed, nor can we know whether MEs/MVs formed before or after a cell lysed. One observation which might be related to this issue comes from *F. johnsoniae* where tubes with regular diameters were seen stemming mainly from cells with a noticeably wavy OM (45 examples), while pearling tubes and OMVs stemmed primarily from cells with a smooth outer membrane (> 100 examples). Compare, for example, the cells in Figures 1e and S6 and Movies S2 and S6 (wavy OM) to those in Figures 3d, 7a and f (smooth OM).

In *C. crescentus*, we observed for the first time “nanopods,” a structure previously reported in *D. acidovorans* [25]. Both of these species are diderms with an S-layer, suggesting that nanopods may be a general form for OMVs in bacteria with this type of cell envelope. Nanopods were proposed to help disperse OMVs in the partially hydrated environment of the soil where *D. acidovorans* lives; it will be interesting to study their function in aquatic *C. crescentus*.

Examining protein complexes associated with OMEs and OMVs, some seemed to reflect a continuation of the same complexes found on the membrane from which the extensions stemmed, such as the T4aP basal body in *M. xanthus* [40]. Others, however, were only observed on MEs and not on cells. This could be because the complexes are related to the formation of the MEs, or it might simply reflect the fact that these extensions are generally thinner and less crowded than the bacterial periplasm, making the complexes easier to see in cryo-tomograms. Interestingly, the crown-like complex we observed in *M. xanthus*, *P. aeruginosa* and *P. flexibilis* was exclusively associated with the membranes of lysed cells; we never observed it on OMEs and OMVs stemming from intact cells in *M. xanthus*. We observed a morphologically-similar crown-like structure with different dimensions in purified naturally-shed MEs/MVs of *S. oneidensis*, where we cannot know whether they arose from intact or lysed cells. The crown-like structures are remarkably large and their function remains a mystery. Due to the disruption of membranes in lysed cells, the topology of these complexes is difficult to unravel. However, these structures share a morphological similarity to a membrane-associated dome protein complex recently described on the limiting membrane of the lamellar bodies inside alveolar cells [41].

Similarly, regarding the different, trapezoidal structure in *S. oneidensis*, the fact that it was seen on both the outside and inside of purified MVs suggests that some of the purified vesicles adopted an inside-out orientation during purification (a documented phenomenon [42]). Interestingly, the overall architecture and dimensions of this trapezoidal structure are reminiscent of those of a recently-solved structure of the *E. coli* polysaccharide co-polymerase WzzB [43]. We hope future investigation by methods like mass spectrometry will characterize these novel ME/MV-associated protein complexes.

In *F. johnsoniae*, we observed secretin-like particles at the tip of ∼70% of tubes stemming from the OM. This strong spatial correlation suggests a role for the secretin-like complex in the formation of MEs in this species. Based on homology, the GspD-like T2SS secretin is a strong candidate for the complex. Interestingly, though, we did not identify any secretin-like (or full T2SS-like) particles in the main cell envelope of *F. johnsoniae* cells. While we could have missed them in the denser periplasm compared to the less-crowded OMEs and OMVs, it is possible that the structures are specifically associated with the formation of OMEs in this species. As these MEs stem only from the OM, there is no IM- embedded energy source for the complex, suggesting that they are not functional secretion systems and raising the question of what function they may serve. It is possible that the OMVs and OMEs form to dispense of the secretin.

These complexes also indicate that MEs/MVs may provide an ideal system to investigate membrane-embedded structures in their native environment at higher resolution. For example, it remains unclear how secretins of various secretion systems are situated within the outer membrane. All high-resolution structures were detergent solubilized, and most *in situ* structures have low resolution due to cell thickness [44]. Purifying *F. johnsoniae* OMVs and performing high- resolution subtomogram averaging on the secretin-like complex might shed light on this question.

Early in the history of life, lipid vesicles and elementary protocells likely experienced destabilizing conditions such as repeated cycles of dehydration and rehydration [45]. The binding of prebiotic amino acids to lipid vesicles can help stabilize them in such conditions [46] and it is conceivable that with billions of years of evolution, variations of these stabilized lipid structures acquired roles that conferred fitness advantages on bacterial species in various environments. Today, the ability of bacteria to extend their membranes to form tubes or vesicles is a widespread phenomenon with many important biological functions. We hope that the structural classification we present here will serve as a helpful reference for future studies in this growing field.

## Supporting information

**Figure S1:**
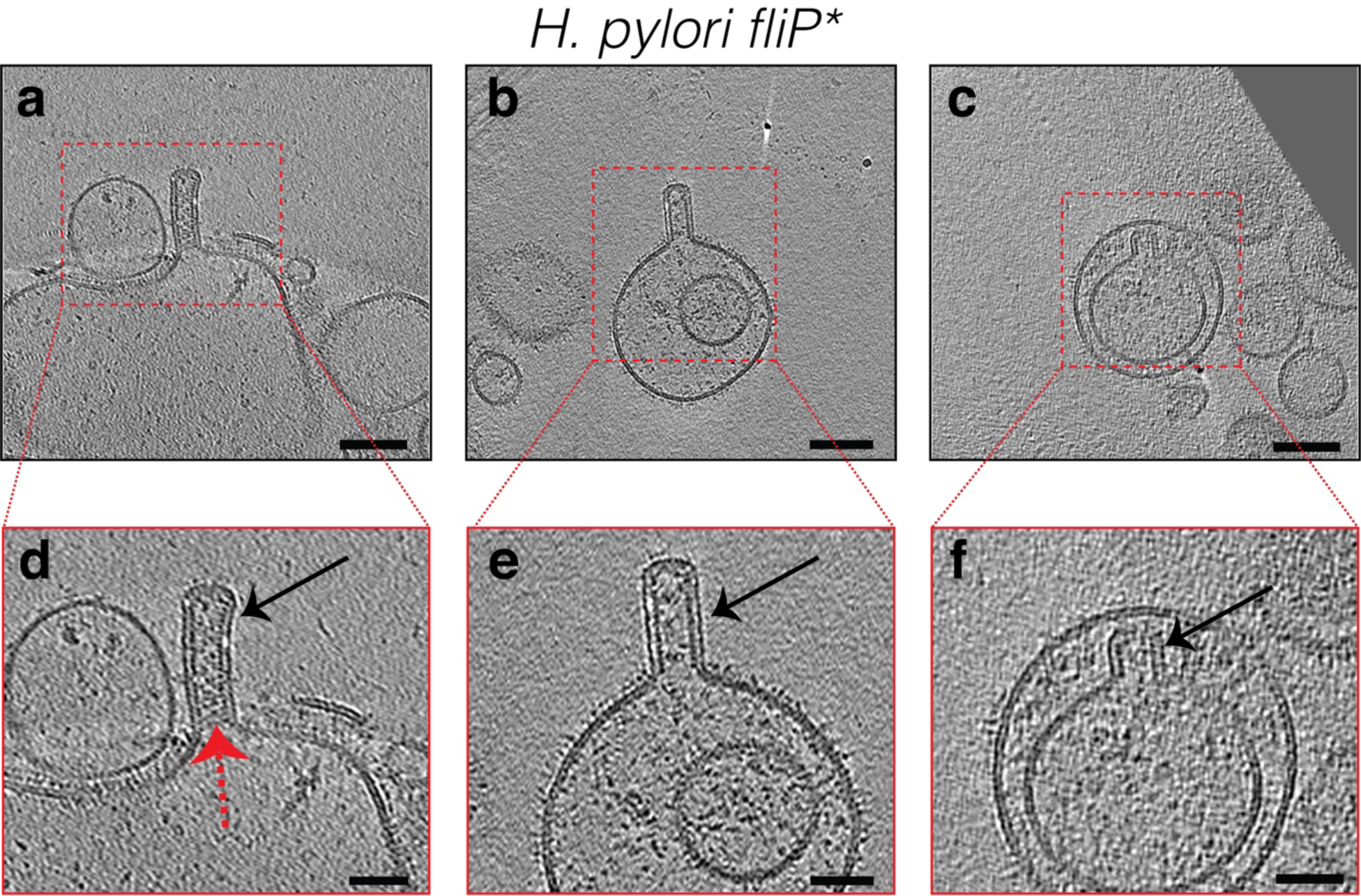
Slices through electron cryo-tomograms of lysed *H. pylori* cells illustrating the presence of OM tubes in vesicles resulting from cell lysis (black arrows). Dashed red arrow in (d) points to the scaffold structure inside the tube. Scale bars are 100 nm in (a-c) and 50 nm in (d-f).

**Figure S2:**
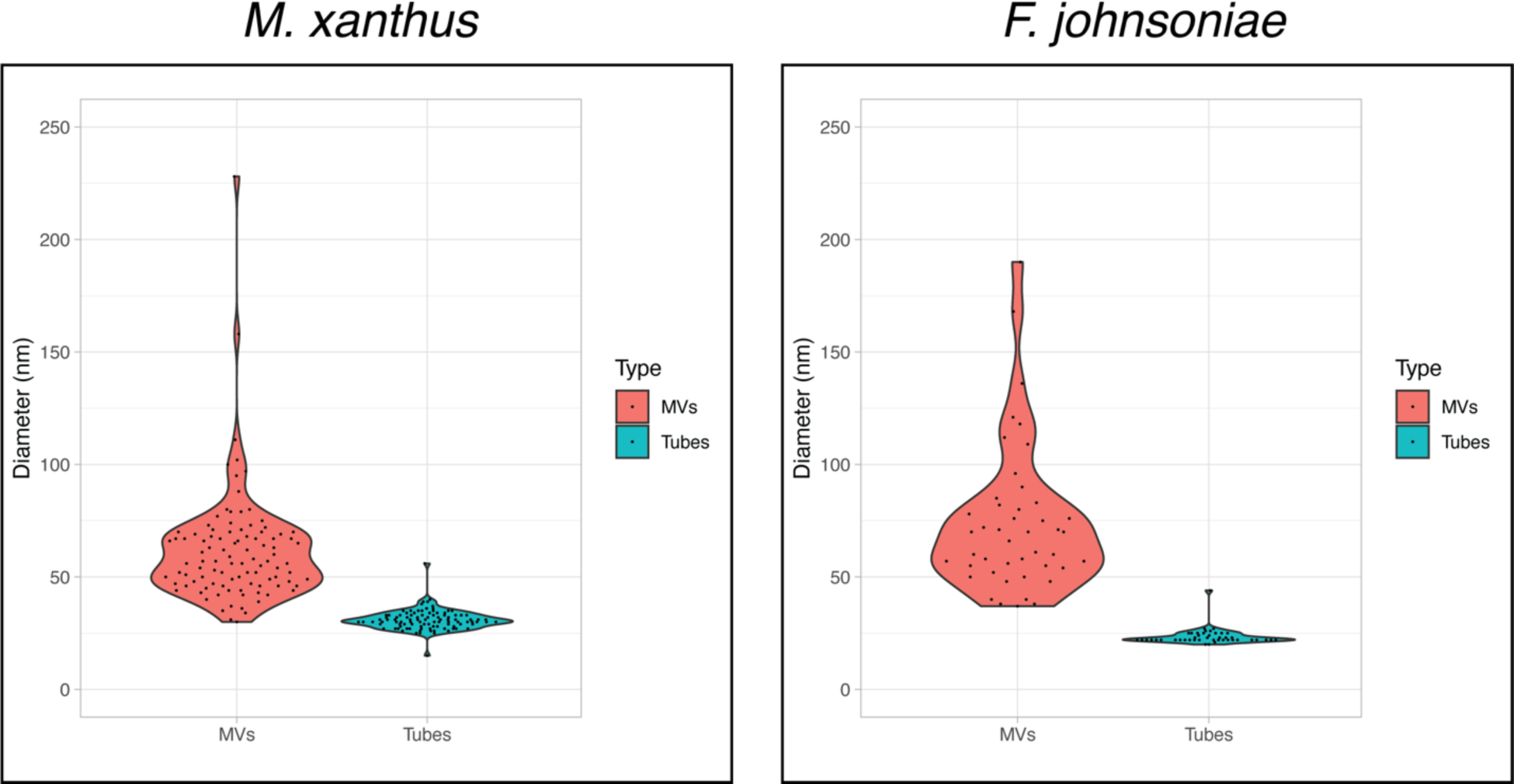
Violin plots of the sizes of OMVs and OM tubes in *M. xanthus* (100 randomly picked- examples of each) and *F. johnsoniae* (45 randomly-picked examples of each). For both species, p<0.001 (determined using t-Test: Two-sample assuming unequal variances).

**Figure S3:**
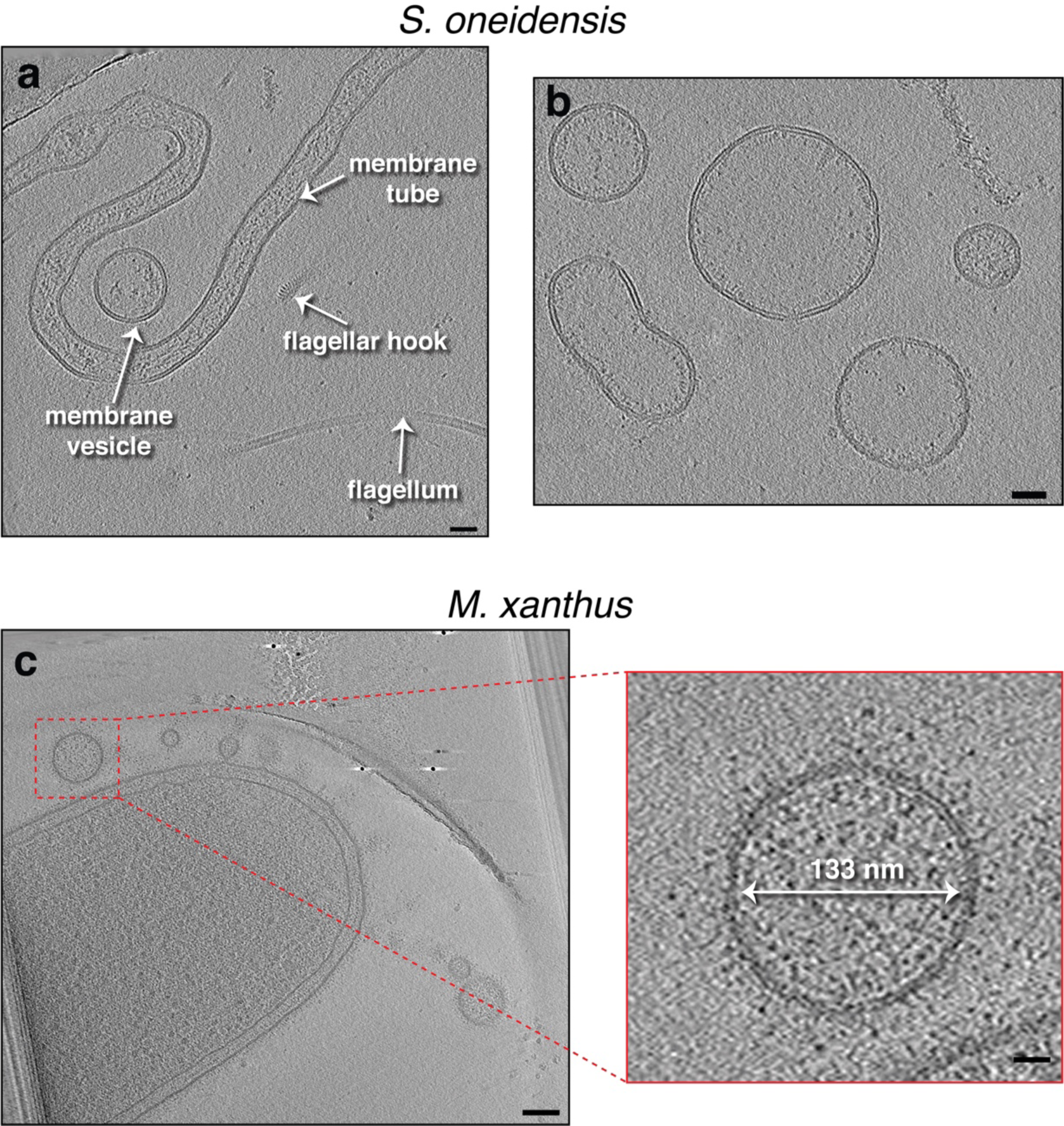
a & b) Slices through electron cryo-tomograms of purified MEs and MVs from *S. oneidensis*. Scale bars are 10 nm. **c)** A slice through an electron cryo-tomogram of an *M. xanthus* cell highlighting the presence of OMVs. Scale bar is 100 nm, and 20 nm in the enlargement on the right.

**Figure S4:**
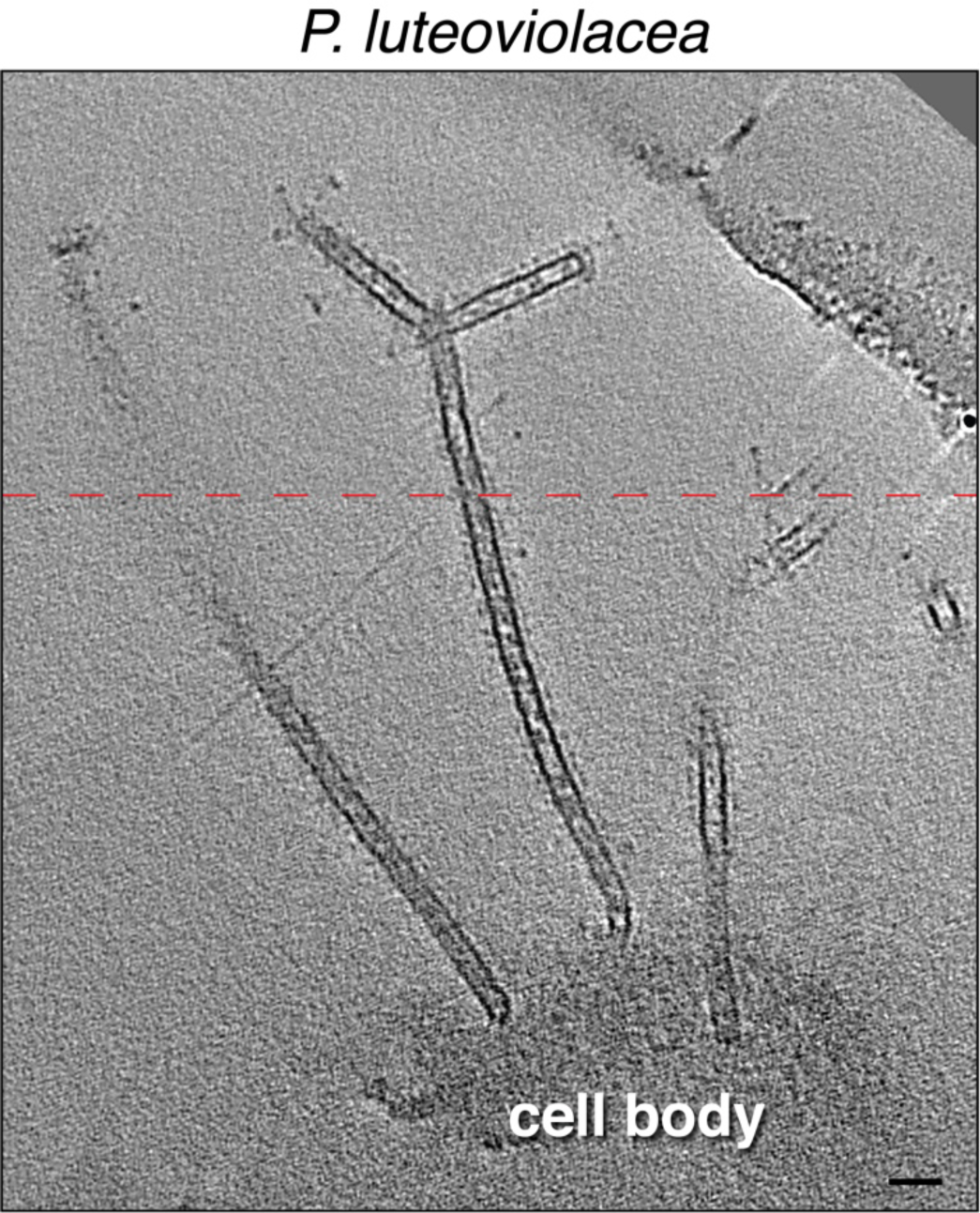
A slice through an electron cryo-tomogram of a lysed *P. luteoviolacea* cell illustrating a bifurcated 20-nm wide membrane tube. Scale bar is 50 nm. Dashed red line indicates a composite image of two slices through the tomogram at different z-heights.

**Figure S5:**
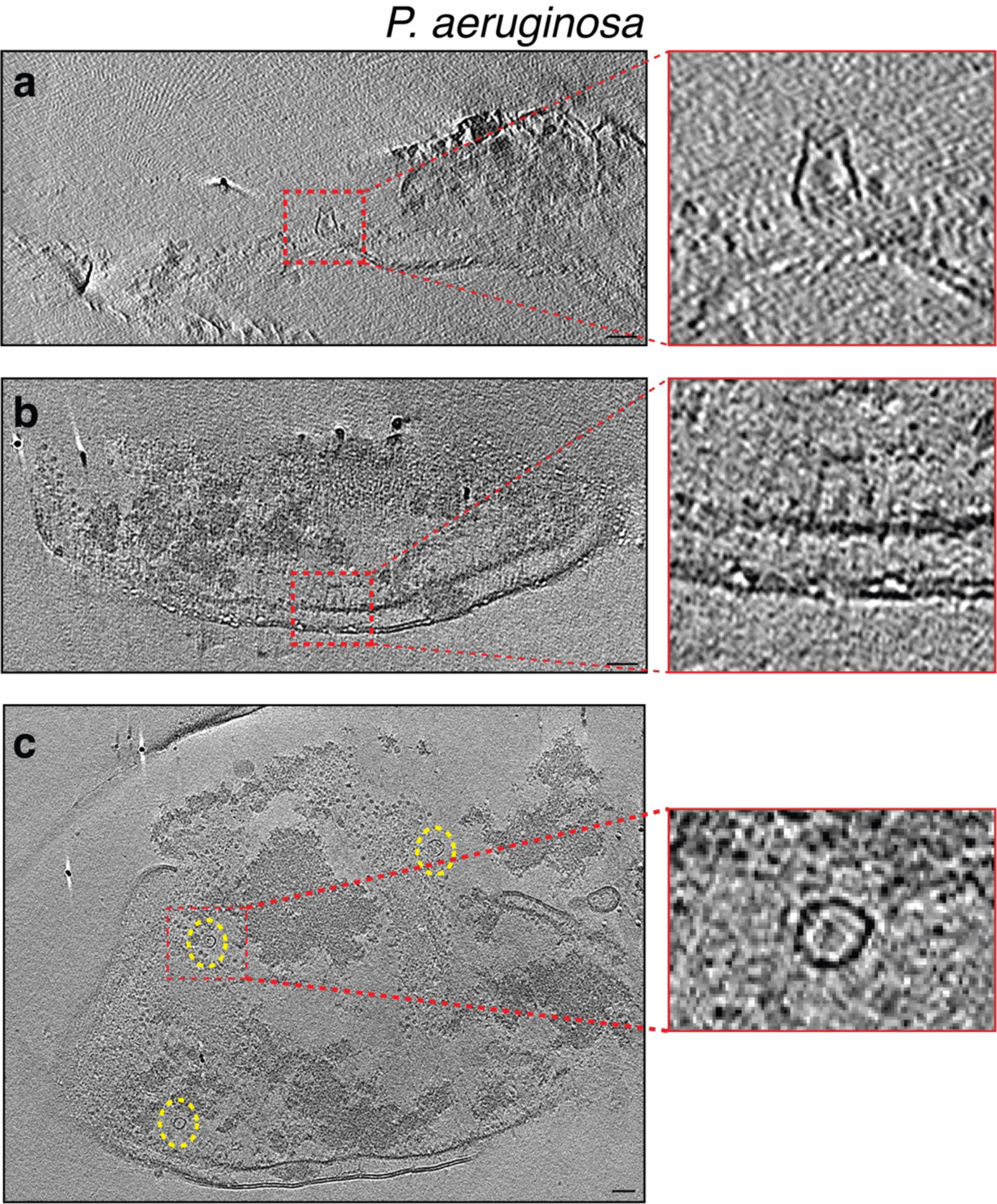
Slices through electron cryo-tomograms of lysed *P. aeruginosa* cells indicating the presence of crown-like structures in side views (**a & b**) and top view (**c**, dashed yellow ellipses). Panels on the right are enlargements of the boxed areas. Scale bars are 50 nm.

**Figure S6:**
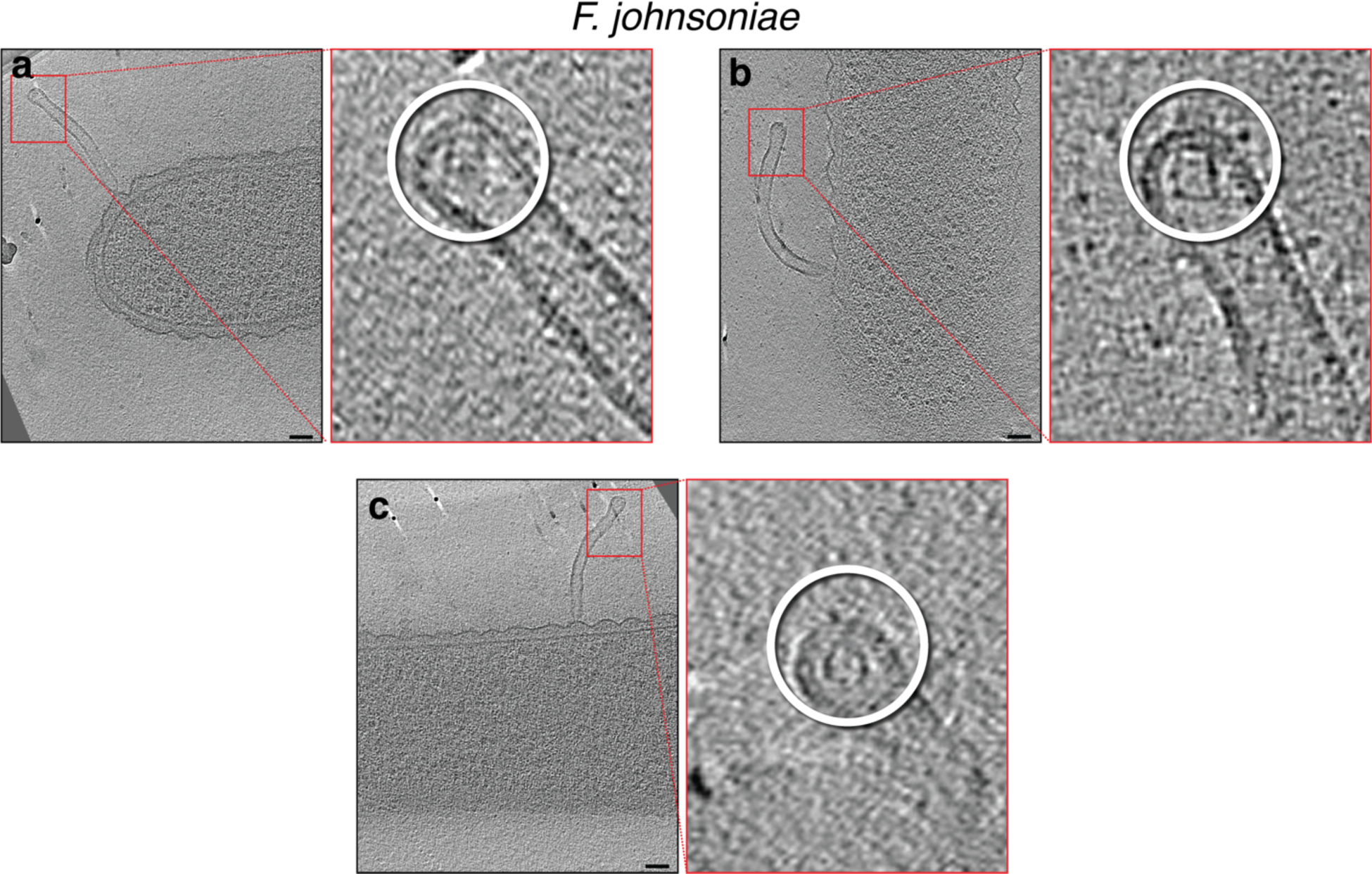
Slices through electron cryo-tomograms of *F. johnsoniae* (with wavy OM) illustrating tubes stemming from cells with secretin-like complexes at their tips, as highlighted in the enlargements on the right (white circles). Note that the rotation of the slices on the left is optimized to show the full tube stemming from the cell, while the rotation of the enlargements on the right is optimized to show the best view of the secretin-like complex. Scale bars are 50 nm.

**Figure S7:**
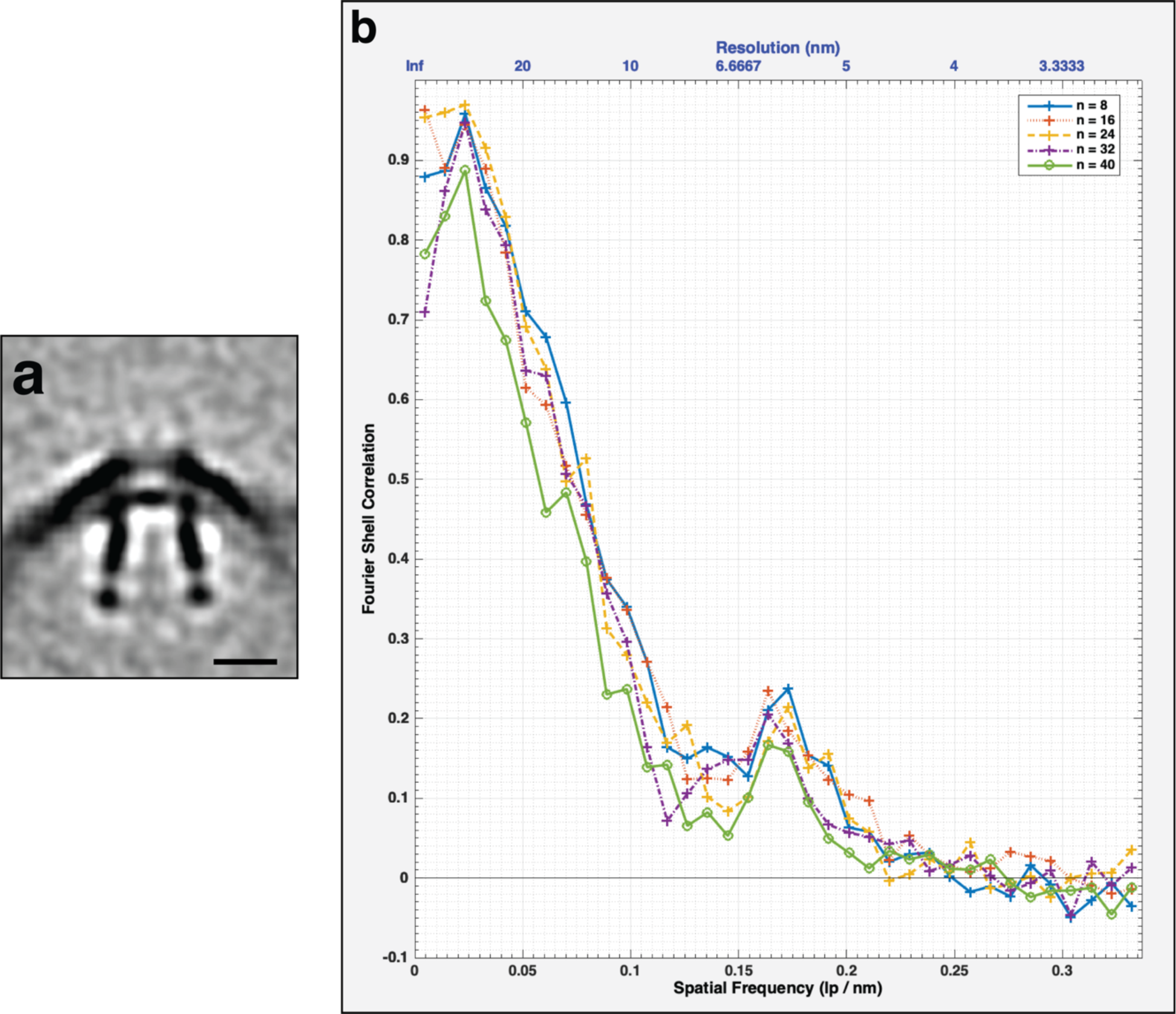
a) Central slice through the unsymmetrized subtomogram average of the secretin-like complex present in OM extensions in *F. johnsoniae*. Scale bar is 10 nm. **b)** FSC curve of the subtomogram average shown in (a). The different colored curves represent different subsets of particles.

## Supplementary movies

**Movie S1:** An electron cryo-tomogram of an *M. xanthus* cell with multiple outer membrane tubes stemming from the cell.

**Movie S2:** An electron cryo-tomogram of an *F. johnsoniae* cell with outer membrane tubes stemming from the cell. Note the wavy outer membrane of the cell.

**Movie S3:** An electron cryo-tomogram of an *M. xanthus* cell with a pearling outer membrane tube stemming from the cell.

**Movie S4:** An electron cryo-tomogram of an *M. xanthus* cell with multiple branched outer membrane tubes stemming from the cell.

**Movie S5:** An electron cryo-tomogram of a *C. crescentus* cell with a nanopod (black arrow) close to the cell.

**Movie S6:** An electron cryo-tomogram of an *F. johnsoniae* cell highlighting the presence of secretin-like particles at the tips of outer membrane tubes.

## Materials and Methods

### Strains and growth conditions

*H. gracilis* cells were grown as described in reference [47]. *P. luteoviolacea* were grown as described in reference [48]. *M. magneticum* were grown as described in reference [49]. *P. flexibilis* 706570 were grown in lactose growth medium. *Shewanella oneidensis* MR-1 cells were grown, as detailed in reference [50], in Luria Bertani (LB) media under aerobic conditions at 30°C with shaking at 200 rpm until they reached OD600 of ∼3. *Myxococcus xanthus* PilY1.3-sfGFP, *M. xanthus* Δ*tsaP*, and *M. xanthus* SA6892 strains were grown as described in reference [40]. *Borrelia burgdorferi* B31 ATCC 35210 and *Helicobacter hepaticus* ATCC 51449 cells were grown in standard media (see reference [51] and references therein).

*Caulobacter crescentus* was cultured in M2G and M2 media (prepared as described in reference [52].) 5 mL of M2G were inoculated with a frozen stock of *C. crescentus* NA 1000 and grown overnight at 28 ℃. 5 mL of the overnight culture was diluted in 15 mL M2G and grown at 28 ℃ with a shaking speed of 200 rpm for ∼2 hours until mid-log phase (OD600 0.4-0.5). The sample was then centrifuged at 5200 ⅹ g for 6 minutes at 4 ℃ (same temperature for all subsequent centrifugation steps) and the pellet was resuspended in 1 mL M2 solution. The resuspended cells were transferred into a 2-mL microcentrifuge tube and centrifuged at 5200 ⅹ g for 5 minutes. All but ∼250 μL of supernatant was removed, 650 μL M2 was added and the pellet was resuspended. 900 μL cold Percoll (Sigma Aldrich) was added and the sample was centrifuged at 15,000 ⅹ g for 20 minutes. Samples were taken from the bottom of the tube to select swarmer cells.

Cells of *Flavobacterium johnsoniae* strain CJ2618 (a wild-type strain overexpressing FtsZ, ATCC 17061) were taken from a glycerol stock, streaked onto a CYE plate with 10ug/mL tetracycline and grown at 25 °C. Subsequently, 5 mL of motility medium (MM) was inoculated with colonies from the plate and the culture was incubated at 25° C with 80 rpm shaking overnight. Then another 5 mL MM was inoculated with 80 uL of starter culture and placed at 25° C with no shaking until the next day when the cells were harvested and prepared for plunge-freezing.

*Helicobacter pylori* mutants (Δ*fliM fliP\**, Δ*fliO fliP\**, Δ*flgS fliP\**, Δ*fliG fliP\**, Δ*fliQ fliP\**) were grown from glycerol stocks on sheep blood agar at 37 °C with 5% CO2 for 48 hours and then either plunge-frozen directly or the cells were spread on another plate and left to grow for 24 hours before plunge-freezing. No difference could be discerned between the two samples by cryo-ET.

*Flavobacterium anhuiense* (strain 98, see reference [53]) and *Chitinophaga pinensis* (strain 94, see reference [53]) cells were grown overnight in 1/10 TSB at 25°C and 300 rpm shaking in 50 ml cultures. For sample preparation, cells were first concentrated by centrifugation. 3 μL aliquots of the cell suspension were applied to glow-discharged R2/2 200 mesh copper Quantifoil grids (Quantifoil Micro Tools), the sample was pre-blotted for 30 seconds, and then blotted for 2.5 seconds (*Flavobacterium anhuiense*) and 1 second (*Chitinophaga pinensis*). Grids were pre- blotted and blotted at 20 °C and at 95 % humidity. Subsequently, the grids were plunge-frozen in liquid ethane using an automated Leica EM GP system (Leica Microsystems) and stored in liquid nitrogen.

### Purification of Shewanella oneidensis OMVs

*S. oneidensis* OMVs were purified as described in reference [50]. First, *S. oneidensis* were grown in LB media until they reached OD600 of 3. Subsequently, the cells were centrifuged at 5000 x g for 20 minutes at 4°C; the pellet contained whole cells while the supernatant contained the OMVs. To remove any cells present in the supernatant, it was filtered through a 0.45 µm filter. Subsequently, the supernatant was centrifuged at 38,400 x g for one hour at 4°C; the OMVs were in the resultant pellet. The pellet was resuspended in 20 ml of 50 mM HEPES pH 6.8 buffer, filtered through a 0.22 µm filter, spun again as described above and ultimately resuspended in 50 mM HEPES pH 6.8.

### Cryo-ET sample preparation and imaging

For cellular samples, 10 nm gold beads were first coated with BSA (bovine serum albumin) and then mixed with the cells. Subsequently, 4 µl of this mixture was applied to a glow-discharged, thick carbon-coated, R2/2, 200 mesh copper Quantifoil grid (Quantifoil Micro Tools) in an FEI Vitrobot chamber with 100% humidity. Excess fluid was blotted away with filter paper and the grid was plunge-frozen in a mixture of ethane/propane. For the purified OMVs of *S. oneidensis*, the sample was first diluted to a 0.4 mg/ml concentration before it was applied to the grid [50]. Cryo-ET imaging of the samples was done either on an FEI Polara 300-keV field emission gun transmission electron microscope equipped with a Gatan imaging filter and a K2 Summit direct electron detector in counting mode, or a Thermo Fisher Titan Krios 300-keV field emission gun transmission electron microscope equipped with a Gatan imaging filter and a K2 Summit counting electron detector. For data collection, either the UCSF Tomography [54] or SerialEM [55] software was used. For OMVs, tilt-series spanned -60° to 60° with an increment of 3°, an underfocus of 1- 5 µm, and a cumulative electron dose of 121 e/Å^2^. For *F. johnsoniae*, tilt-series spanned -55° to 55° with 1° increment, an underfocus of 4 µm, a cumulative electron dose of 100 e/Å^2^, and a 3.9 Å pixel size. For *M. xanthus,* tilt-series spanned -60° to 60° with an increment of 1°, an underfocus of 6 µm, and a cumulative electron dose of 180 e/Å^2^. For *B. burgdorferi*, tilt-series spanned -60° to 60° with 1° increment, an underfocus of 10 µm, and a cumulative electron dose of 160 e/Å^2^. For *H. hepaticus*, tilt-series spanned -60° to 60° with increments of 1°, an underfocus of 12 µm, and a cumulative electron dose of 165 e/Å^2^.

*Flavobacterium anhuiense* and *Chitinophaga pinensis* images were recorded with a Gatan K3 Summit direct electron detector equipped with a Gatan GIF Quantum energy filter with a slit width of 20 eV. Images were taken at magnification corresponding to a pixel size of 3.28 Å (*Chitinophaga pinensis*) and 4.4 Å (*Flavobacterium anhuiense*). Tilt series were collected using SerialEM with a bidirectional dose-symmetric tilt scheme (-60° to 60°, starting from 0°) with a 2° increment. The defocus was set to – 8- 10 μm and the cumulative exposure per tilt series was 100 e-/A^2^. Images were reconstructed with the IMOD software package.

### Image processing and subtomogram averaging

Reconstruction of tomograms of cellular samples was done using the automatic RAPTOR pipeline implemented in the Jensen laboratory at Caltech [31]. Tomograms of purified *S. oneidensis* OMVs were reconstructed using a combination of ctffind4 [56] and the IMOD software package [57]. Subtomogram averaging was done using the PEET program [58], with 2-fold symmetry applied along the particle y-axis.

**Table S1:** A summary of the species included in this study and the major membrane structures identified in each species.

| Species | Class | Features Observed |  |  |  |  |  |  |  |
| --- | --- | --- | --- | --- | --- | --- | --- | --- | --- |
|  |  | Tubes |  |  |  | Vesicle chains |  | Budding/<br>vesicles | Nanopods |
|  |  | Uniform<br>diameter<br>-<br>scaffold | Uniform<br>diameter<br>- no<br>scaffold | Variable<br>diameter | Pearling | Connectors | No<br>connectors |  |  |
| *Shewanella oneidensis* | Gammaproteobacteria |  |  |  |  |  | see Ref [6] | >100 |  |
| *Pseudoalteromonas luteoviolacea* | Gammaproteobacteria |  | ~100 |  | ~10 |  |  |  |  |
| *Hylemonella gracilis* | Betaproteobacteria |  |  | 3 | 4 |  |  | 15 |  |
| *Delftia acidovorans* | Betaproteobacteria |  |  |  |  |  |  |  | see Ref [25] |
| *Magnetospirillum magneticum* | Alphaproteobacteria |  |  |  |  |  |  | 49 |  |
| *Caulobacter crescentus* | Alphaproteobacteria |  |  |  |  |  |  |  | 53 |
| *Helicobacter hepaticus* | Epsilonproteobacteria |  |  |  | 2 |  |  |  |  |
| *Helicobacter pylori* | Epsilonproteobacteria | >100 |  |  |  |  |  |  |  |
| *Myxococcus xanthus* | Deltaproteobacteria |  | >100 |  | >100 |  |  | >100 |  |
| *Borrelia burgdorferi* | Spirochaetes |  | 9 |  |  | 19 |  | 16 |  |
| *Flavobacterium johnsoniae* | Flavobacteria |  | ~45 |  | ~15 |  |  | >100 |  |
| *Flavobacterium anhuiense* | Flavobacteria |  |  | 5 | 7 |  | 4 | >100<br>(including the<br>teardrop-like<br>extensions) |  |
| *Chitinophaga pinensis* | Chitinophagia |  |  | 11 | 12 |  | 3 | 81<br>(including the<br>teardrop-like<br>extensions) |  |

## Acknowledgements

This project was funded by the NIH (grant R35 GM122588 to G.J.J) and a Baxter postdoctoral fellowship from Caltech to M.K. Cryo-ET work was done in the Beckman Institute Resource Center for Transmission Electron Microscopy at the California Institute of Technology. We are grateful to Prof. Martin Pilhofer for collecting the *P. luteoviolacea* data and for critically reading the manuscript. We thank Prof. Elitza I Tocheva for collecting the *D. acidovorans* data. We thank Prof. Mohamed El-Naggar for insights into preparing *S. oneidensis* samples and Dr. Yuxi Liu for discussions. Briegel lab data was collected at the Netherlands Center for Electron Nanoscopy with support from Dr. Wen Yang. This data was collected with support from the National Roadmap for Large-Scale Research Infrastructure 2017 – 2018 with project number 184.034.014, which is financed in part by the Dutch Research Council (NWO). This work was also supported by the NWO OCENW.GROOT.2019.063 grant.

## References

1. Schwechheimer C, Kuehn MJ. Outer-membrane vesicles from Gram-negative bacteria: biogenesis and functions. Nature Reviews Microbiology. 2015;13: 605–619. doi:10.1038/nrmicro3525

2. Jan AT. Outer Membrane Vesicles (OMVs) of Gram-negative Bacteria: A Perspective Update. Frontiers in Microbiology. 2017;8. doi:10.3389/fmicb.2017.01053

3. D’Souza G, Shitut S, Preussger D, Yousif G, Waschina S, Kost C. Ecology and evolution of metabolic cross-feeding interactions in bacteria. Natural Product Reports. 2018;35: 455– 488. doi:10.1039/C8NP00009C

4. Toyofuku M, Nomura N, Eberl L. Types and origins of bacterial membrane vesicles. Nature Reviews Microbiology. 2019;17: 13–24. doi:10.1038/s41579-018-0112-2

5. Pirbadian S, Barchinger SE, Leung KM, Byun HS, Jangir Y, Bouhenni RA, et al. Shewanella oneidensis MR-1 nanowires are outer membrane and periplasmic extensions of the extracellular electron transport components. Proceedings of the National Academy of Sciences. 2014;111: 12883–12888. doi:10.1073/pnas.1410551111

6. Subramanian P, Pirbadian S, El-Naggar MY, Jensen GJ. Ultrastructure of *Shewanella oneidensis* MR-1 nanowires revealed by electron cryotomography. Proceedings of the National Academy of Sciences. 2018;115: E3246–E3255. doi:10.1073/pnas.1718810115

7. Ducret A, Fleuchot B, Bergam P, Mignot T. Direct live imaging of cell–cell protein transfer by transient outer membrane fusion in Myxococcus xanthus. eLife. 2013;2. doi:10.7554/eLife.00868

8. Wei X, Vassallo CN, Pathak DT, Wall D. Myxobacteria Produce Outer Membrane-Enclosed Tubes in Unstructured Environments. Journal of Bacteriology. 2014;196: 1807– 1814. doi:10.1128/JB.00850-13

9. Remis JP, Wei D, Gorur A, Zemla M, Haraga J, Allen S, et al. Bacterial social networks: structure and composition of *Myxococcus xanthus* outer membrane vesicle chains: Membrane vesicle chains and membrane network of *M. xanthus*. Environmental Microbiology. 2014;16: 598–610. doi:10.1111/1462-2920.12187

10. Reyes-Robles T, Dillard RS, Cairns LS, Silva-Valenzuela CA, Housman M, Ali A, et al. *Vibrio cholerae* Outer Membrane Vesicles Inhibit Bacteriophage Infection. DiRita VJ, editor. Journal of Bacteriology. 2018;200. doi:10.1128/JB.00792-17

11. Fischer T, Schorb M, Reintjes G, Kolovou A, Santarella-Mellwig R, Markert S, et al. Biopearling of Interconnected Outer Membrane Vesicle Chains by a Marine Flavobacterium. Cann I, editor. Applied and Environmental Microbiology. 2019;85. doi:10.1128/AEM.00829-19

12. Møller J, Barnes A, Dalsgaard I, Ellis A. Characterisation of surface blebbing and membrane vesicles produced by Flavobacterium psychrophilum. Diseases of Aquatic Organisms. 2005;64: 201–209. doi:10.3354/dao064201

13. McCaig WD, Koller A, Thanassi DG. Production of Outer Membrane Vesicles and Outer Membrane Tubes by Francisella novicida. Journal of Bacteriology. 2013;195: 1120–1132. doi:10.1128/JB.02007-12

14. Laanto E, Penttinen RK, Bamford JK, Sundberg L-R. Comparing the different morphotypes of a fish pathogen - implications for key virulence factors in Flavobacterium columnare. BMC Microbiology. 2014;14: 170. doi:10.1186/1471-2180-14-170

15. Chang Y-W, Shaffer CL, Rettberg LA, Ghosal D, Jensen GJ. In Vivo Structures of the Helicobacter pylori cag Type IV Secretion System. Cell Reports. 2018;23: 673–681. doi:10.1016/j.celrep.2018.03.085

16. Hampton CM, Guerrero-Ferreira RC, Storms RE, Taylor JV, Yi H, Gulig PA, et al. The Opportunistic Pathogen Vibrio vulnificus Produces Outer Membrane Vesicles in a Spatially Distinct Manner Related to Capsular Polysaccharide. Frontiers in Microbiology. 2017;8. doi:10.3389/fmicb.2017.02177

17. Brown L, Wolf JM, Prados-Rosales R, Casadevall A. Through the wall: extracellular vesicles in Gram-positive bacteria, mycobacteria and fungi. Nature Reviews Microbiology. 2015;13: 620–630. doi:10.1038/nrmicro3480

18. Bhattacharya S, Baidya AK, Pal RR, Mamou G, Gatt YE, Margalit H, et al. A Ubiquitous Platform for Bacterial Nanotube Biogenesis. Cell Reports. 2019;27: 334–342.e10. doi:10.1016/j.celrep.2019.02.055

19. Dubey GP, Ben-Yehuda S. Intercellular Nanotubes Mediate Bacterial Communication. Cell. 2011;144: 590–600. doi:10.1016/j.cell.2011.01.015

20. Baidya AK, Bhattacharya S, Dubey GP, Mamou G, Ben-Yehuda S. Bacterial nanotubes: a conduit for intercellular molecular trade. Current Opinion in Microbiology. 2018;42: 1–6. doi:10.1016/j.mib.2017.08.006

21. Pande S, Shitut S, Freund L, Westermann M, Bertels F, Colesie C, et al. Metabolic cross- feeding via intercellular nanotubes among bacteria. Nature Communications. 2015;6. doi:10.1038/ncomms7238

22. Benomar S, Ranava D, Cárdenas ML, Trably E, Rafrafi Y, Ducret A, et al. Nutritional stress induces exchange of cell material and energetic coupling between bacterial species. Nat Commun. 2015;6: 6283. doi:10.1038/ncomms7283

23. Baidya AK, Rosenshine I, Ben-Yehuda S. Donor-delivered cell wall hydrolases facilitate nanotube penetration into recipient bacteria. Nat Commun. 2020;11: 1938. doi:10.1038/s41467-020-15605-1

24. Pal RR, Baidya AK, Mamou G, Bhattacharya S, Socol Y, Kobi S, et al. Pathogenic E. coli Extracts Nutrients from Infected Host Cells Utilizing Injectisome Components. Cell. 2019;177: 683–696.e18. doi:10.1016/j.cell.2019.02.022

25. Shetty A, Chen S, Tocheva EI, Jensen GJ, Hickey WJ. Nanopods: A New Bacterial Structure and Mechanism for Deployment of Outer Membrane Vesicles. Hensel M, editor. PLoS ONE. 2011;6: e20725. doi:10.1371/journal.pone.0020725

26. Marguet E, Gaudin M, Gauliard E, Fourquaux I, le Blond du Plouy S, Matsui I, et al. Membrane vesicles, nanopods and/or nanotubes produced by hyperthermophilic archaea of the genus Thermococcus. Biochemical Society Transactions. 2013;41: 436–442. doi:10.1042/BST20120293

27. Bar-Ziv R, Moses E. Instability and “Pearling” States Produced in Tubular Membranes by Competition of Curvature and Tension. Physical Review Letters. 1994;73: 1392–1395. doi:10.1103/PhysRevLett.73.1392

28. Pospíšil J, Vítovská D, Kofroňová O, Muchová K, Šanderová H, Hubálek M, et al. Bacterial nanotubes as a manifestation of cell death. Nature Communications. 2020;11. doi:10.1038/s41467-020-18800-2

29. Bos J, Cisneros LH, Mazel D. Real-time tracking of bacterial membrane vesicles reveals enhanced membrane traffic upon antibiotic exposure. Science Advances. 2021;7: eabd1033. doi:10.1126/sciadv.abd1033

30. Sivabalasarma S, Wetzel H, Nußbaum P, van der Does C, Beeby M, Albers S-V. Analysis of Cell–Cell Bridges in Haloferax volcanii Using Electron Cryo-Tomography Reveal a Continuous Cytoplasm and S-Layer. Front Microbiol. 2021;11: 612239. doi:10.3389/fmicb.2020.612239

31. Ding HJ, Oikonomou CM, Jensen GJ. The Caltech Tomography Database and Automatic Processing Pipeline. Journal of Structural Biology. 2015;192: 279–286. doi:10.1016/j.jsb.2015.06.016

32. Ortega DR, Oikonomou CM, Ding HJ, Rees-Lee P, Alexandria, Jensen GJ. ETDB-Caltech: A blockchain-based distributed public database for electron tomography. Promponas VJ, editor. PLOS ONE. 2019;14: e0215531. doi:10.1371/journal.pone.0215531

33. Fukumura T, Makino F, Dietsche T, Kinoshita M, Kato T, Wagner S, et al. Assembly and stoichiometry of the core structure of the bacterial flagellar type III export gate complex. Stock A, editor. PLOS Biology. 2017;15: e2002281. doi:10.1371/journal.pbio.2002281

34. Fabiani FD, Renault TT, Peters B, Dietsche T, Gálvez EJC, Guse A, et al. A flagellum- specific chaperone facilitates assembly of the core type III export apparatus of the bacterial flagellum. Stock A, editor. PLOS Biology. 2017;15: e2002267. doi:10.1371/journal.pbio.2002267

35. Minamino T, Kawamoto A, Kinoshita M, Namba K. Molecular Organization and Assembly of the Export Apparatus of Flagellar Type III Secretion Systems. Berlin, Heidelberg: Springer Berlin Heidelberg; 2019. doi:10.1007/82_2019_170

36. Lertsethtakarn P, Ottemann KM, Hendrixson DR. Motility and Chemotaxis in *Campylobacter* and *Helicobacter*. Annual Review of Microbiology. 2011;65: 389–410. doi:10.1146/annurev-micro-090110-102908

37. Wang X, Han Q, Chen G, Zhang W, Liu W. A Putative Type II Secretion System Is Involved in Cellulose Utilization in Cytophaga hutchisonii. Frontiers in Microbiology. 2017;8. doi:10.3389/fmicb.2017.01482

38. Ghosal D, Kim KW, Zheng H, Kaplan M, Truchan HK, Lopez AE, et al. In vivo structure of the Legionella type II secretion system by electron cryotomography. Nature Microbiology. 2019 [cited 22 Nov 2019]. doi:10.1038/s41564-019-0603-6

39. Yuan F, Alimohamadi H, Bakka B, Trementozzi AN, Day KJ, Fawzi NL, et al. Membrane bending by protein phase separation. Proc Natl Acad Sci USA. 2021;118: e2017435118. doi:10.1073/pnas.2017435118

40. Chang Y-W, Rettberg LA, Treuner-Lange A, Iwasa J, Søgaard-Andersen L, Jensen GJ. Architecture of the type IVa pilus machine. Science. 2016;351: aad2001. doi:10.1126/science.aad2001

41. Klein S, Wimmer BH, Winter SL, Kolovou A, Laketa V, Chlanda P. Post-correlation on- lamella cryo-CLEM reveals the membrane architecture of lamellar bodies. Commun Biol. 2021;4: 137. doi:10.1038/s42003-020-01567-z

42. Kaplan M, Narasimhan S, de Heus C, Mance D, van Doorn S, Houben K, et al. EGFR Dynamics Change during Activation in Native Membranes as Revealed by NMR. Cell. 2016;167: 1241–1251.e11. doi:10.1016/j.cell.2016.10.038

43. Wiseman B, Nitharwal RG, Widmalm G, Högbom M. Structure of a full-length bacterial polysaccharide co-polymerase. Nat Commun. 2021;12: 369. doi:10.1038/s41467-020-20579-1

44. Weaver SJ, Ortega DR, Sazinsky MH, Dalia TN, Dalia AB, Jensen GJ. CryoEM structure of the type IVa pilus secretin required for natural competence in Vibrio cholerae. Nature Communications. 2020;11. doi:10.1038/s41467-020-18866-y

45. Damer B, Deamer D. Coupled Phases and Combinatorial Selection in Fluctuating Hydrothermal Pools: A Scenario to Guide Experimental Approaches to the Origin of Cellular Life. Life. 2015;5: 872–887. doi:10.3390/life5010872

46. Cornell CE, Black RA, Xue M, Litz HE, Ramsay A, Gordon M, et al. Prebiotic amino acids bind to and stabilize prebiotic fatty acid membranes. Proceedings of the National Academy of Sciences. 2019;116: 17239–17244. doi:10.1073/pnas.1900275116

47. Kaplan M, Sweredoski MJ, Rodrigues JPGLM, Tocheva EI, Chang Y-W, Ortega DR, et al. Bacterial flagellar motor PL-ring disassembly subcomplexes are widespread and ancient. Proceedings of the National Academy of Sciences. 2020; 201916935. doi:10.1073/pnas.1916935117

48. Shikuma NJ, Pilhofer M, Weiss GL, Hadfield MG, Jensen GJ, Newman DK. Marine Tubeworm Metamorphosis Induced by Arrays of Bacterial Phage Tail-Like Structures. Science. 2014;343: 529–533. doi:10.1126/science.1246794

49. Cornejo E, Subramanian P, Li Z, Jensen GJ, Komeili A. Dynamic Remodeling of the Magnetosome Membrane Is Triggered by the Initiation of Biomineralization. mBio. 2016;7. doi:10.1128/mBio.01898-15

50. Phillips DA, Zacharoff LA, Hampton CM, Chong GW, Malanoski AP, Metskas LA, et al. A Prokaryotic Membrane Sculpting BAR Domain Protein. Microbiology; 2020 Jan. doi:10.1101/2020.01.30.926147

51. Briegel A, Ortega DR, Tocheva EI, Wuichet K, Li Z, Chen S, et al. Universal architecture of bacterial chemoreceptor arrays. Proceedings of the National Academy of Sciences. 2009;106: 17181–17186. doi:10.1073/pnas.0905181106

52. Schrader JM, Shapiro L. Synchronization of Caulobacter Crescentus for Investigation of the Bacterial Cell Cycle. Journal of Visualized Experiments. 2015 [cited 15 Nov 2020]. doi:10.3791/52633

53. Carrión VJ, Perez-Jaramillo J, Cordovez V, Tracanna V, de Hollander M, Ruiz-Buck D, et al. Pathogen-induced activation of disease-suppressive functions in the endophytic root microbiome. Science. 2019;366: 606–612. doi:10.1126/science.aaw9285

54. Zheng SQ, Keszthelyi B, Branlund E, Lyle JM, Braunfeld MB, Sedat JW, et al. UCSF tomography: an integrated software suite for real-time electron microscopic tomographic data collection, alignment, and reconstruction. J Struct Biol. 2007;157: 138–147. doi:10.1016/j.jsb.2006.06.005

55. Mastronarde DN. Automated electron microscope tomography using robust prediction of specimen movements. J Struct Biol. 2005;152: 36–51. doi:10.1016/j.jsb.2005.07.007

56. Rohou A, Grigorieff N. CTFFIND4: Fast and accurate defocus estimation from electron micrographs. Journal of Structural Biology. 2015;192: 216–221. doi:10.1016/j.jsb.2015.08.008

57. Kremer JR, Mastronarde DN, McIntosh JR. Computer visualization of three-dimensional image data using IMOD. J Struct Biol. 1996;116: 71–76. doi:10.1006/jsbi.1996.0013

58. Nicastro D. The Molecular Architecture of Axonemes Revealed by Cryoelectron Tomography. Science. 2006;313: 944–948. doi:10.1126/science.1128618

